# Hybrid asexuality as a primary reproductive barrier: on the interconnection between asexuality and speciation

**DOI:** 10.1101/038299

**Authors:** Karel Janko, Jan Pačes, Hilde Wilkinson-Herbots, Rui J Costa, Jan Roslein, Pavel Drozd, Nataliia Iakovenko, Jakub Rídl, Jan Kočí, Radka Reifová, Věra Šlechtová, Lukáš Choleva

## Abstract

Speciation usually proceeds in a continuum from intensively hybridizing populations until the formation of irreversibly isolated species. Restriction of interspecific gene flow may often be achieved by gradual accumulation of intrinsic postzygotic incompatibilities with hybrid infertility typically evolving more rapidly than inviability. A reconstructed history of speciation in European loaches *(Cobitis)* reveals that accumulation of postzygotic reproductive incompatibilities may take an alternative, in the literature largely neglected, pathway through initiation of hybrids’ asexuality rather than through a decrease in hybrids’ fitness. Combined evidence shows that contemporary *Cobitis* species readily hybridize in hybrid zones, but their gene pools are isolated as hybridization produces infertile males and fertile but clonally reproducing females that cannot mediate introgressions. Nevertheless, coalescent analyses indicated intensive historical gene flow during earlier stages of *Cobitis* diversification, suggesting that non-clonal hybrids must have existed in the past. The revealed patterns imply that during the initial stages of speciation, hybridization between little diverged species produced recombinant hybrids mediating gene flow, but growing divergence among species caused disrupted meiosis in hybrids resulting in their clonality, which acts as a barrier to gene flow. Comparative analysis of published data on other fish hybrids corroborated the generality of our findings; the species pairs producing asexual hybrids were more genetically diverged than those pairs producing fertile sexual hybrids but less diverged than species pairs producing infertile hybrids. Hybrid asexuality therefore appears to evolve at lower divergence than other types of postzygotic barriers and might thus represent a primary reproductive barrier in many taxa.

## Introduction

Speciation may be accomplished by various reproductive isolating mechanisms (RIMs) that restrict interspecific gene flow. These RIMs can be initiated either by divergent ecological or sexual selection that creates prezygotic and extrinsic postzygotic reproductive isolation or by gradual accumulation of genetic incompatibilities that cause intrinsic postzygotic isolation, i.e. hybrid sterility and inviability (rev. in [1]). The study of intrinsic postzygotic isolation has a strong tradition in evolutionary biology [2, 3] and comparative studies showed that hybridization capability decreases as one moves from closely to distantly related pairs of taxa [4–7]. Although the rate at which intrinsic postzygotic RIMs accumulate is probably nonlinear [8–10] and varies among taxa [11], hybrid infertility generally evolves at lower genetic distances than hybrid inviability [5, 12] and according to Haldane’s rule, both hybrid sterility and inviability evolve faster in the heterogametic sex [13].

Here we suggest that speciation driven by intrinsic postzygotic barriers may take an alternative, in literature largely neglected, pathway: the accumulation of genetic incompatibilities may lead to distortion of hybrids’ reproductive mode towards asexuality already at early stages of the species diversification when hybrids’ fitness has not yet considerably decreased. Given that obligate asexuals are generally incapable to introgress their genomes into related sexual species (e.g. [14, 15]), such modification of hybrids’ reproductive mode may effectively block interspecific gene exchange and contribute to speciation even if other forms of RIMs are absent. This hypothesis is consistent with the observation that asexual organisms often appear as hybrids between sexually reproducing species [16]. Various mutually not exclusive theories attempted to explain the association between hybridization and asexuality (rev. in [17]). Among those, it has been proposed early in 20^th^ century that the likelihood of initiation of hybrid asexuality could depend on the genetic distance between hybridizing species [18], possibly because the coexistence of distinct genomes within progenitor egg cells may deregulate the pathways underlying meiosis [19–21]. Moritz et al. [22] formulated the “balance hypothesis” as a candidate general model of the hybridization-asexuality association. It postulates that asexuality can arise only when the genomes of parental species accumulated enough mutational incompatibilities to alter meiosis or gametogenesis in hybrids yet not enough to seriously compromise hybrid viability or fertility.

From this perspective, one could regard hybrid asexuality as a special case of the accumulation of Dobzhansky-Muller incompatibilities that disrupt critical processes - sexual reproduction in this case – and thus directly contributes to speciation. Unfortunately, there is only scarce empirical support for the link between accumulating divergences among hybridizing genomes and the initiation of asexuality [23] and hence, it remains unclear how or even whether hybridization could initiate the production of clonal gametes [17, 24, 25].

This study explores the role of hybrid asexuality in speciation by simultaneous reconstruction of the rates of interspecific gene flow and evolution of asexuality during the diversification history of European spined loaches *(Cobitis)*. These bottom-dwelling fish colonized Europe in several phylogenetic lineages during the Tertiary period. While isolated rivers of Mediterranean Europe are inhabited by diverse *Cobitis* species clustering in deeply divergent phylogenetic lineages *(Cobitis* Lineage I – V *sensu* [26]), vast areas of non-Mediterranean Europe were colonized by a single lineage *(Cobitis* Lineage V) comprising several morphologically and ecologically very similar species. Among these, four closely related species (C. *tanaitica, C. taenia*, *C. taurica* and *C. pontica)* have Ponto Caspian distribution (C. *taenia* further colonised Northern and Western Europe) while their distant relative, *C. elongatoides* is distributed throughout the Danubian basin. Their ranges have been fluctuating over the Quaternary [27] and currently display a parapatric distribution with overlapping zones located in Central Europe, the lower Danube, and Southern Ukraine [28] (Fig 1). Hybridization has been documented to take place in these zones and reproductive contact between *C. elongatoides* and either *C. taenia, C. tanaitica* or *C. pontica* has been documented to result in clonally reproducing all-females hybrid lineages [24, 28, 29]. These hybrid lineages achieved remarkable evolutionary success and colonised most of European continent, some of them having achieved considerable ages (i.e. the so called Hybrid Clades I and II are as ancient as 0.35 and 0.25 Mya, respectively [27, 30]). Clonal reproduction of these hybrid lineages theoretically prevents any interspecific gene flow but we have recently documented evidence for intensive gene flow between *C. elongatoides* and *C. tanaitica* leading to the fixation of an introgressed *C*. *elongatoides*-like mitochondrion over the entire *C. tanaitica* distribution range [31].

**Fig 1.**
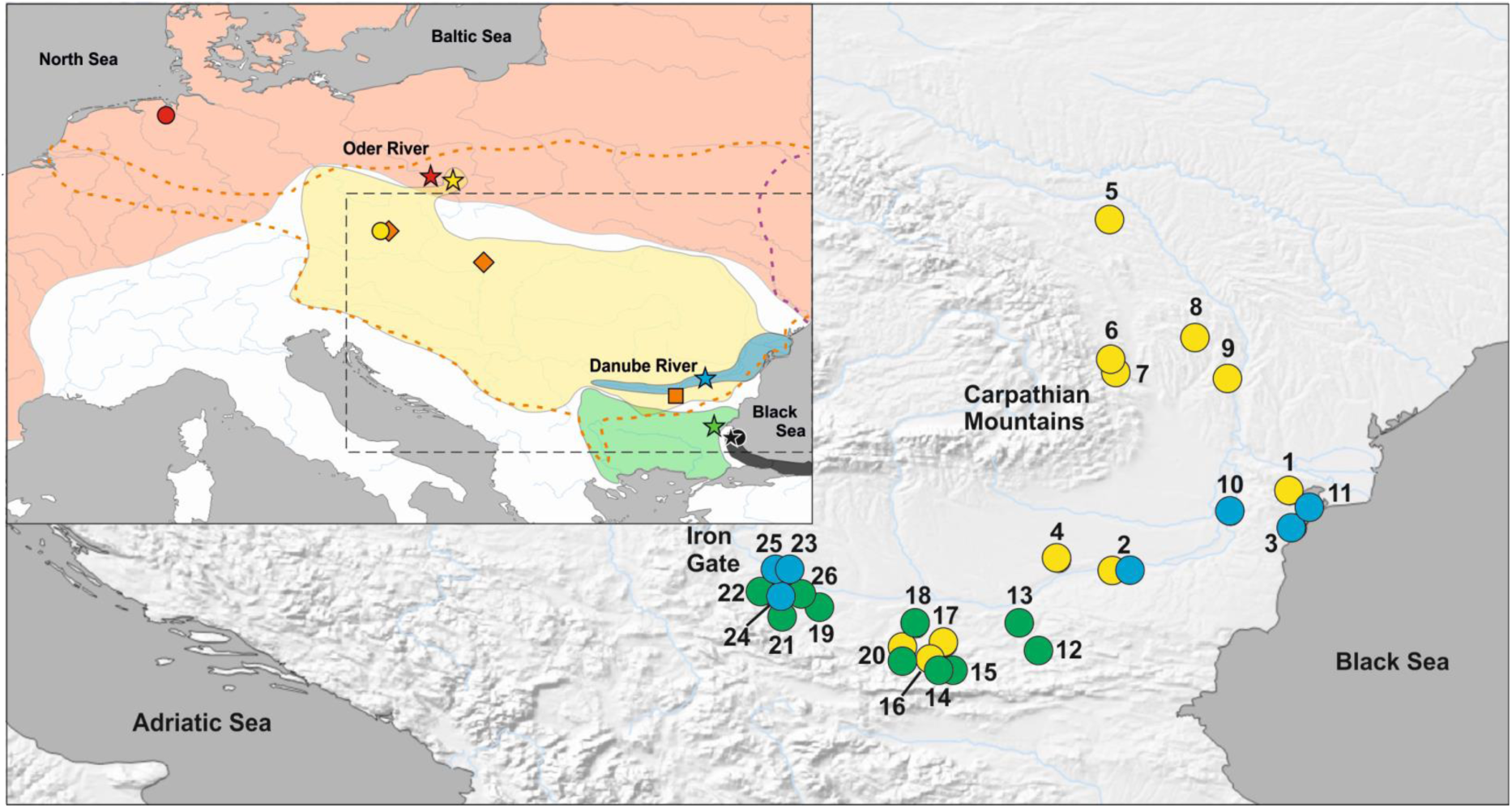
Map of sampling sites and distribution of species and hybrid biotypes.

The inset shows European distribution of studied sexual species. Red stands for *C. taenia* distribution range, yellow for *C. elongatoides*, blue for *C. tanaitica*, black for *C. pontica*, and green for *C. strumicae*. Orange dotted line delimits the distribution of the ancient clonal lineage, the so called Hybrid Clade I, purple dotted line the distribution of the so called Hybrid Clade II. Stars indicate sampling sites of individuals used for transcriptome analyses, circles, squares and diamonds of those used in crossing experiments (circles indicate sampled sexual species, the orange square stands for diploid and diamonds for triploid *C. elongatoides-tanaitica* hybrids used in crossings). Main map shows in detail the Lower Danubian hybrid zone: Yellow circles indicate localities with *C. elongatoides* samples, blue represent *C. tanaitica* and green indicate *C. strumicae*. Locality numbers correspond with Table 2.

**Table 2.**
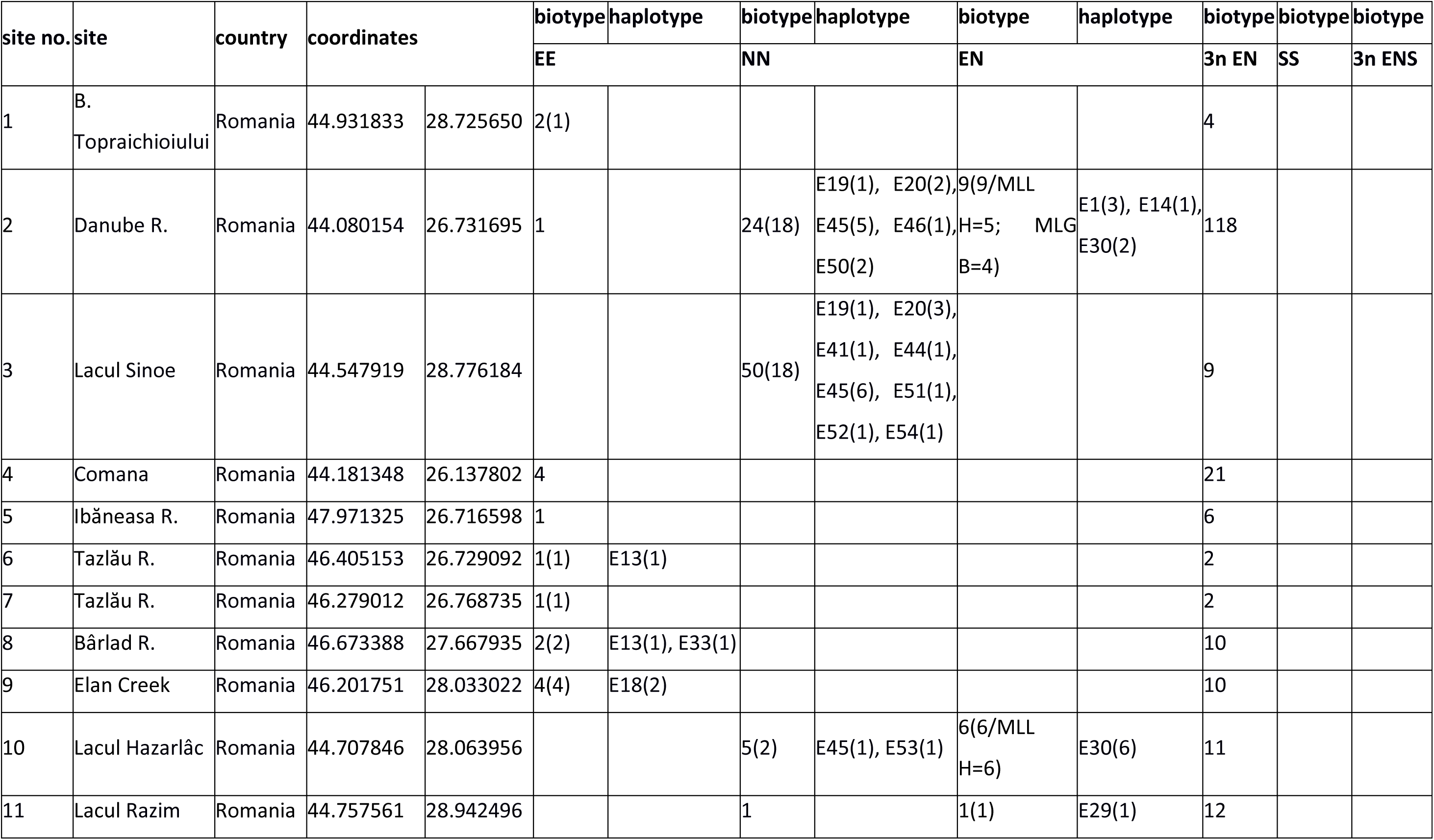
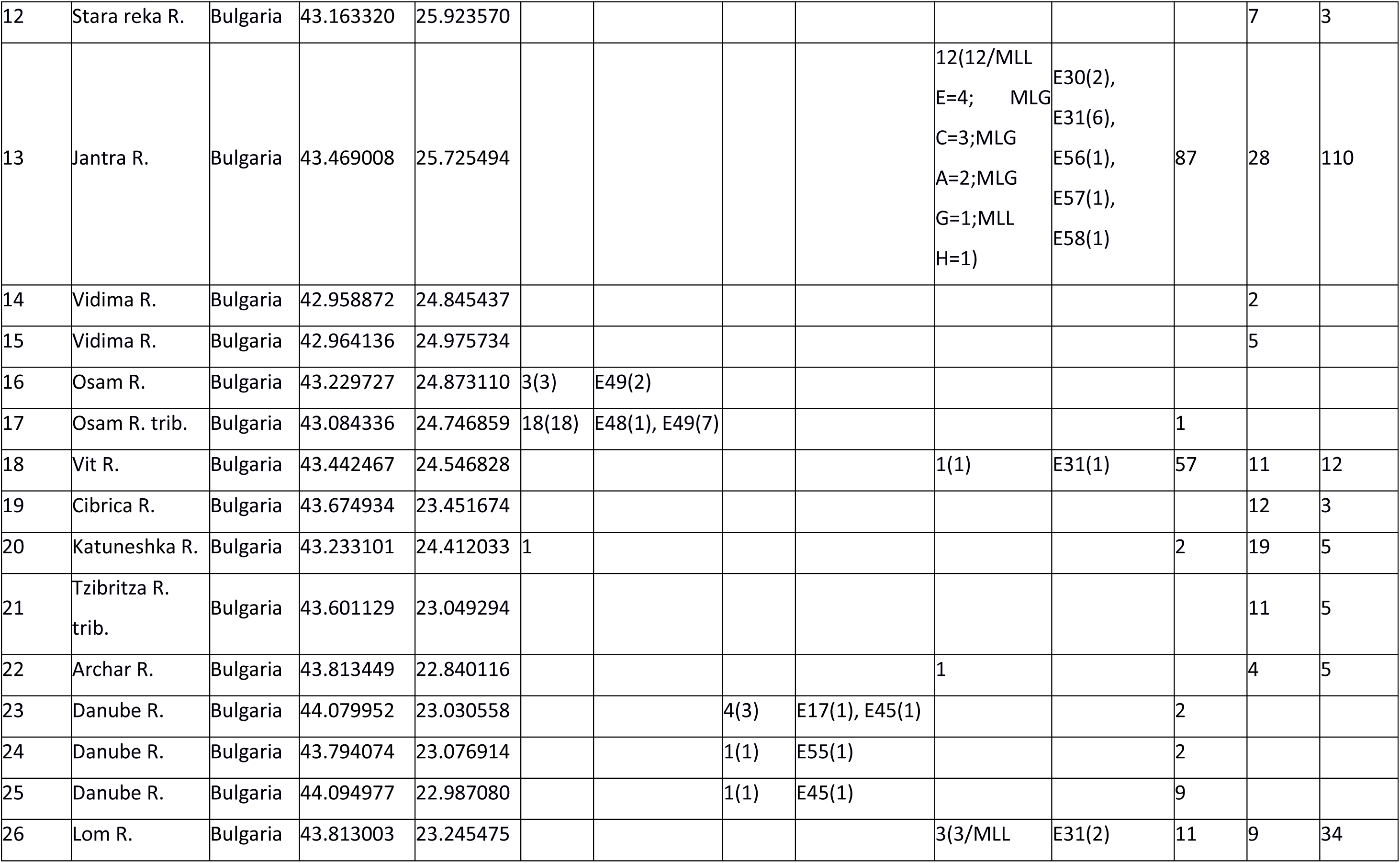
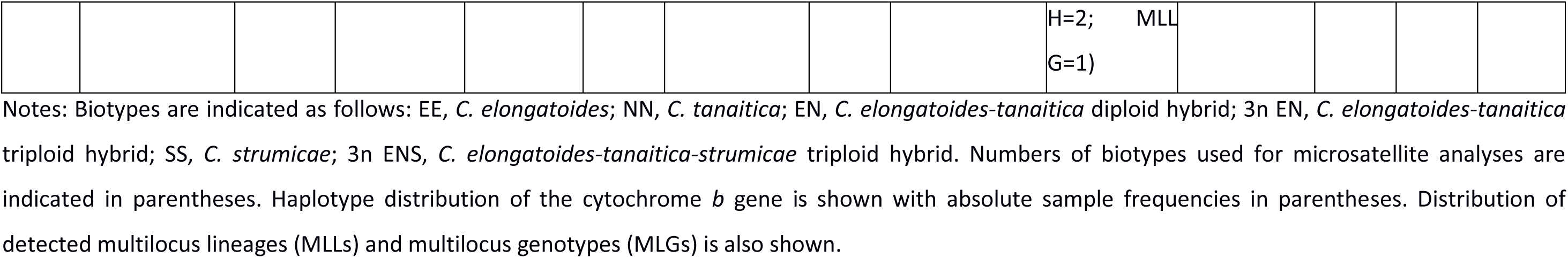
Locality information from the Danube River hybrid zone.

To understand the speciation history in spined loaches and more generally the role of hybrid asexuality in fish speciation, we employed multiple complementary approaches examining a) whether the establishment of reproductive barriers can be primarily accomplished by the formation of hybrid asexuality, b) whether the formation of asexual hybrids does depend on genetic distance between parental species. First, we analyzed the reproductive modes in hybrids between several differently related *Cobitis* species to reveal their asexuality and the extent of the currently observed reproductive isolation. Second, we performed a detailed population genetic analysis of hybrid zones to test if there is any ongoing introgressive hybridization. Third, we employed coalescent analyses of transcriptomic data to estimate levels and timing of historical gene flow among species. The combined evidence implies that the diversification of *Cobitis* species has been accompanied by decreasing intensity of introgressive hybridization. However such a restriction in gene flow has not been accomplished by classical RIMs but rather by the production of asexual hybrids suggesting that asexual hybrids may constitute a primary reproductive barrier between nascent species.

## Results

To test whether the formation of asexual hybrids does depend on genetic distance between parental species and how the formation of hybrid asexuality affects the levels of reproductive isolation we performed the following four analyses.

### Analysis of reproductive modes of interspecific Cobitis hybrids

Firstly, we tested whether *Cobitis* hybrids can mediate interspecific gene flow by analysing reproductive modes of natural and artificial hybrids between several differently related *Cobitis* species.

*1. Reproductive modes of natural C. elongatoides-tanaitica hybrids:* We successfully analysed five backcrossed families stemming from four natural triploid EEN and one diploid EN hybrid females (letters E and N stand for haploid *elongatoides* and *tanaitica* genomes, respectively). Backcross progeny consistently expressed all maternal alleles, suggesting the production of unreduced gametes and lack of segregation. In one allele, single progeny differed from the maternal allele by a single repeat, indicating a mutation event. A part of the progeny also contained a haploid set of paternal alleles indicating that the sperm’s genome is sometimes incorporated, leading to a ploidy increase (Table 1 and S1). We also compared allozyme profiles of eggs and somatic tissues of six EEN females, in order to test for hemiclonal reproduction (i.e., hybridogenesis). Hybridogenesis was supposed to lead to reduced, albeit non-segregating gametes [32–34], which consistently express allozyme products of one parental species only. Nevertheless, we found that allozyme profiles of all eggs were identical to the somatic tissues of their maternal individuals, further suggesting that natural hybrids do not exclude any genome (Table S2).

**Table 1.** Crossing experiments.

| female parent | biotype | origin | male parent | biotype | origin | progeny |  |  | Family ID |
| --- | --- | --- | --- | --- | --- | --- | --- | --- | --- |
|  |  |  |  |  |  | sexual | clonal | polyploid |  |
| EENF1 | EEN | Okna R., Slovakia | EEM1 | EE | Okna R., Slovakia | 0 | 6 | 1 | No. 401 |
| EENF24 | EEN | Okna R., Slovakia | EEM8 | EE | Okna R., Slovakia | 0 | 16 | 0 | No. 424 |
| EENF25 | EEN | Okna R., Slovakia | EEM8 | EE | Okna R., Slovakia | 0 | 9 | 0 | No. 425 |
| EENF10 | EEN | Ipeľ R., Slovakia | EEM9 | EE | Nová Říše, Czech R. | 0 | 9 | 0 | No. 413 |
| 097CENNF1 | EN | Jantra R., Bulgaria | 09EXPM7C | PP | Veleka R., Bulgaria | 0 | 6 | 4 | No. 8 |
| 10EXF1F9C08 | TP | laboratory hybrid | 10EXF1M9C08 | TP | laboratory hybrid | 11 | 6 | 9 | No. 1 |
| F1TFPM062 | TP | laboratory hybrid | F1TFPM065 | TP | laboratory hybrid | 6 | 0 | 0 | No. 17; clutch A |
| F1TFPM062 | TP | laboratory hybrid | F1TFPM066 | TP | laboratory hybrid | 1 | 0 | 0 | No. 17; clutch B |

Notes: E, haploid *C. elongatoides* genome; N, haploid *C. tanaitica* genome; P, haploid *C. pontica* genome; T, haploid *C. taenia* genome; For each family, we indicate different types of progeny: “sexual” denotes a number of progeny obtained from segregating gametes; “clonal”, a number of progeny obtained from clonal gametes; “polyploid”, a number of progeny obtained from fertilized clonal gametes. Note that several clutches from different F1 individuals occurred in the experimental family No. 17.

*2. Reproductive modes of artificial F1 hybrids:* In addition to *C. elongatoides – C. taenia* crosses published in [24], we newly obtained two *C. taenia – C. pontica* primary crosses. Family No. 1 (Tables 1 and S1) produced a number of F1 hybrids, of which one F1 x F1 couple successfully spawned and produced F2 progeny. Second reproducing family (No. 17; Tables 1 and S1) reached maturity in a number of four males and three females in a single aquarium. From this family we obtained two clutches of F2 progeny spawned by single F1 female, which mated with two different F1 males. Both hybrid sexes were viable and fertile in both families No. 1 and No. 17, as evidenced by successful production of F2 progeny. F2 progeny mostly possessed one allele from the mother and the other from the father. Such inheritance patterns suggest the sexual reproduction of C. *taenia-pontica* hybrids in both families. However, family No. 1 also contained different types of F2 progeny: 15 individuals contained the complete set of maternal alleles and 9 of them also possessed the haploid set of paternal alleles (Table 1). Such patterns indicate that C. *taenia-pontica* hybrid females produced not only recombinant sexual gametes but partly also unreduced gametes. The unreduced eggs either developed into a clonal progeny via gynogenesis or into triploid progeny after a true fertilization with a haploid sperm.

### Analysis of C. elongatoides – C. tanaitica hybrid zone and test of ongoing introgressive hybridization

Second, we tested for ongoing gene flow between species by performing population genetic analysis of *C. elongatoides – C. tanaitica* hybrid zone similar to *C. elongatoides – C. taenia* analysed in [30]. 818 *Cobitis* specimens were captured and identified on 26 localities all over the lower Danubian River Basin. Due to morphological similarity, the taxonomical diagnosis was performed by allozyme analysis of diagnostic loci [28], which revealed the presence of *C. elongatoides* at 11 sites and *C. tanaitica* at seven sites. 10 localities were inhabited by distantly related species, *C. strumicae*, belonging to the subgenus Bicanestrinia (lineage III sensu [26]) (Fig 1 and Table 2). Apart from sexual species, we found various hybrid biotypes, which were mostly polyploid but we also encountered 32 diploid hybrids heterozygotic for species-specific alleles at all diagnostic allozyme loci (Table 2).

Microsatellite analysis was successfully performed on 105 diploid individuals. Most diploids had unique combinations of alleles (Table S3), but 28 diploids clustered in eight groups of identical MultiLocus Genotypes (MLG A – MLG H). Such groups of identical individuals were considered as clone mates due to negligible probability of identical genotypes arising from independent sexual events (*p* < 10^-5^). Furthermore, we noticed that two clusters of individuals or MLGs were separated by genetic distances, which were significantly lower than would be expected from independent sexual events (*p* < 0.01). As in [30], both such groups were assigned as MultiLocus Lineages (MLL) and individuals clustering within such lineages were considered as members of the same clone. Altogether, 29 diploid individuals could be assumed to form eight clonal lineages. Allozymes indicated a hybrid state of all such clonal individuals. No mtDNA haplotypes were shared by *C. elongatoides* and *C. tanaitica* (Fig 2). One clone (MLG B) possessed a haplotype E1, which was shared with *C. elongatoides*, while the remaining individuals assigned as hybrids possessed haplotypes clustering in the old Hybrid clade I defined in [27].

**Figure 2.**
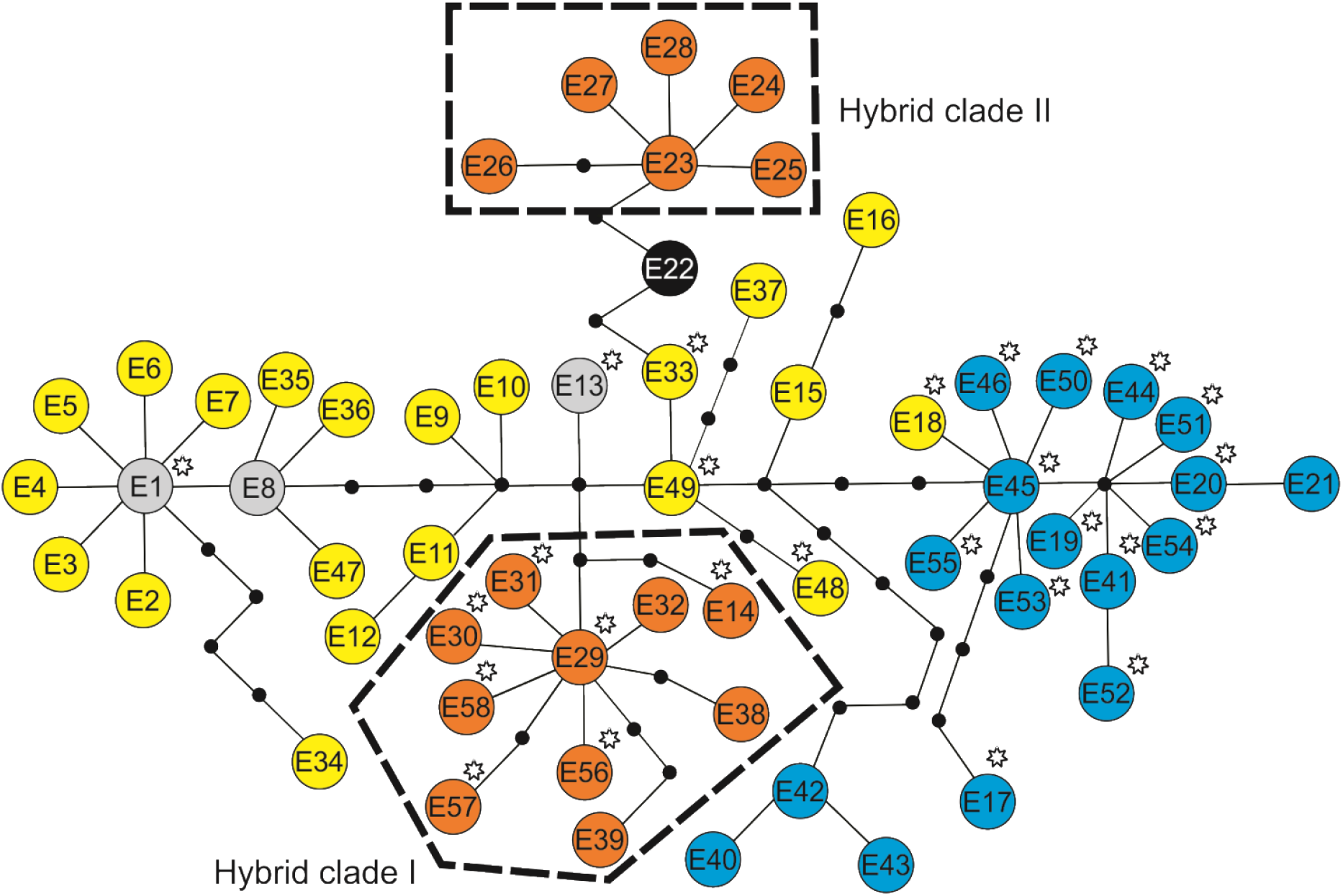
Median-joining haplotype network showing phylogenetic relationships among *C. elongatoides-like* haplotypes of the cytochrome *b* gene.

The network was constructed from previously published haplotypes and those from the current study (with asterisk). Yellow colour denotes haplotypes sampled in *C. elongatoides;* blue in *C. tanaitica;* black in *C. pontica;* orange in *C. elongatoides-tanaitica* hybrid (Hybrid clade I) and *C. elongatoides-taenia* hybrid (Hybrid clade II). Light grey circles denote haplotypes shared by both *C. elongatoides* and hybrids. Small black circles represent missing (unobserved) haplotypes.

Analysis of the combined microsatellite and allozyme data by the Structure provided the likelihood values that converged during the runs and results did not notably change between replicates. Regardless of the locality, the optimal number of clusters *K =* 2 was selected as the bestfit partitioning of the diploid dataset. Altogether, we found three types of diploid individuals. First, we identified those individuals, where the parameter *q* ranged between 0.993 and 0.999 (presumably *C. elongatoides* individuals). Second, we identified those individuals with *q* values between 0.002 and 0.011 (43 presumably *C. tanaitica* individuals). Finally, all 32 individuals identified as hybrids by allozymes had intermediate *q* values (0.554 – 0.783; Fig 3A).

**Figure 3.**
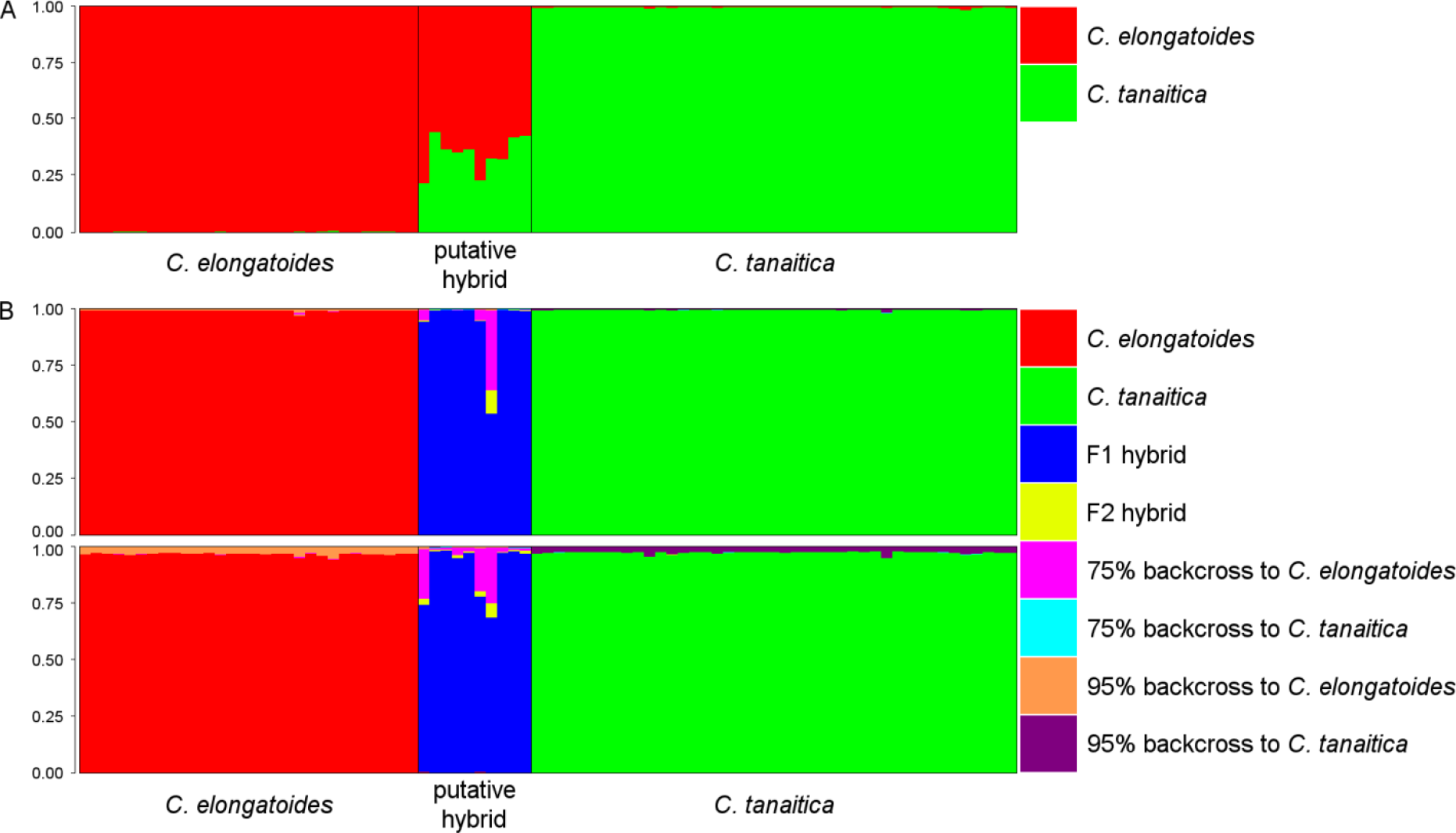
Population genetic analyses of the hybrid zone.

(A) Individual proportion of membership to one of the two species-specific clusters according to Structure for *K* = 2. Each vertical bar represents one individual and colours show proportion of their assignment to respective clusters corresponding to sexual species. For the visual guidance, the individuals are grouped into a priori defined biotypes according to diagnostic allozyme markers (horizontal axis). (B) Classification of individual’s genotype according to NewHybrids. Each vertical bar represents one individual. Each colour represents the posterior probability of an individual belonging to one of the eight different genotypic classes. Individuals are sorted as in (A). Upper pane represents the results with Jeffreys prior and lower pane with the uniform prior.

Data analysis using the NewHybrids software was slightly sensitive to the type of prior applied to *θ* but not to *π*. However, consistent with the Structure, all specimens presumed to be *C. elongatoides* and *C. tanaitica* were always assigned as the pure parental species *(p >* 0.95). Under Jeffrey’s prior, the NewHybrids assigned all but one of the abovementioned hybrids into the F1 class with the probability exceeding 95 %; however, it could not decide in the case of MLL E between F1 (p = 0.53), F2 (p = 0.10), and B1 (p = 0.35) states (Fig 3B). Under the uniform prior, most hybrids were assigned into F1 class, but NewHybrids could not decide between F1 and B1 states of MLL H, MLL E and individual 09BG19K22 (Fig 3B and Table S3).

### Estimation of levels and timing of historical gene flow among the species using transcriptome data

Thirdly, we estimated levels and timing of historical gene flow among the species by analysed single nucleotide polymorphisms (SNP) variability of *C. elongatoides, C. taenia*, *C. tanaitica* and *C. pontica* transcriptomes. As an outgroup we used *C. strumicae* – a species belonging to subgenus *Bicanestrinia* whose divergence from the ingroup was set to 17.7 Mya according to [35], which was sampled in isolated Black Sea tributaries outside of the distribution ranges of ingroup species in order to avoid any possible reproductive contact. The assembly of mRNA from five *C. taenia* specimens comprised 20,385 contigs (potential mRNAs) and was used as a reference for mapping of reads from all species (two individuals of each ingroup species and one outgroup). In total, we identified 187,205 SNPs (Appendix S4), which were then analysed by fitting eight coalescent models assuming different scenarios of species divergence and connectivity [36–40]. The models are characterized in Fig 4 and the results of analyses are described in Table 3. Because the available models only allow fitting pairs of taxa, we separately analysed six datasets corresponding to all pairwise combinations of the four analysed species. Most models were successfully fitted to all datasets but two models did not converge in the case of the *C. taenia – C. tanaitica* dataset.

**Figure 4.**
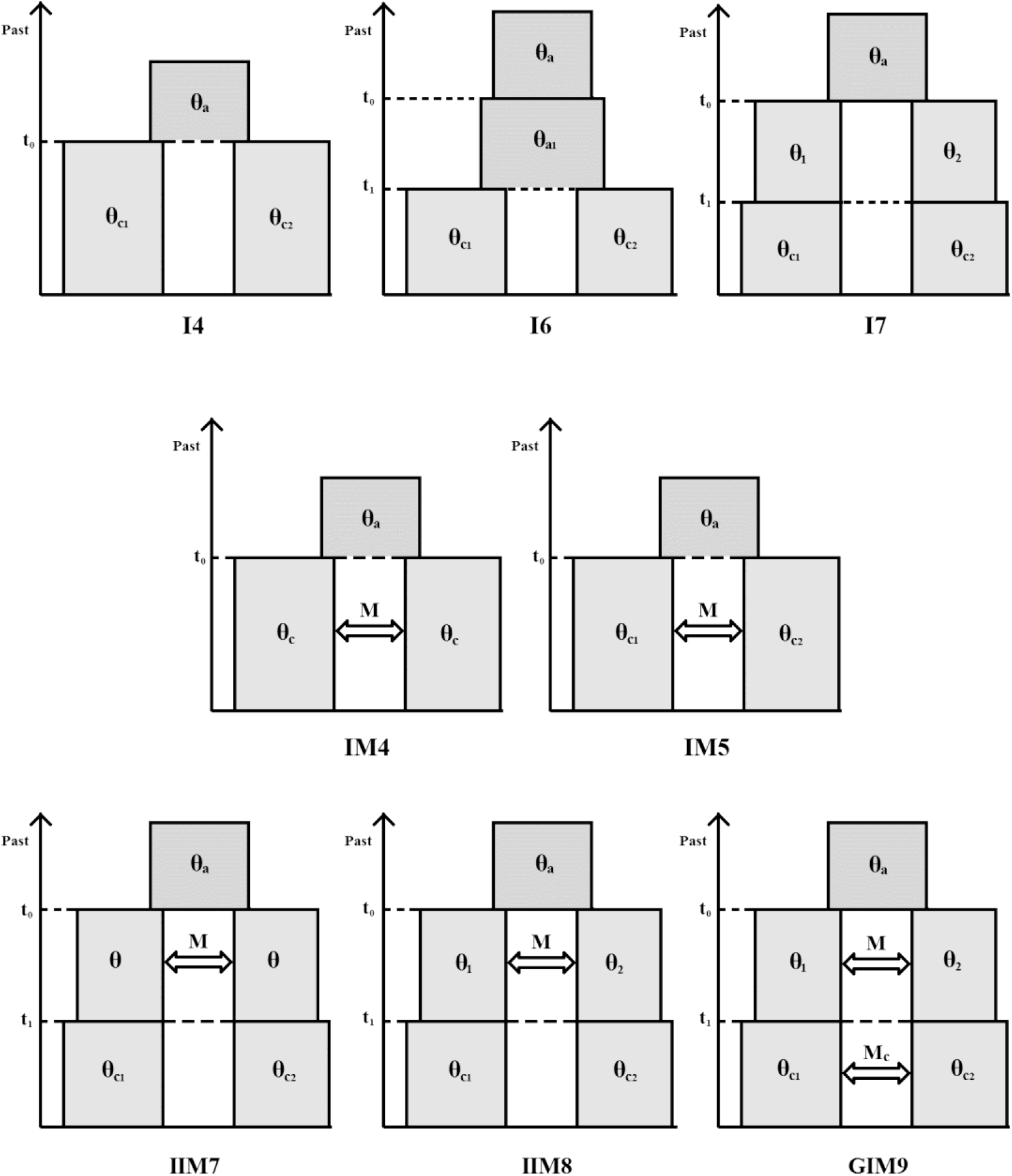
Schematic view of the eight coalescent models.

**Table 3.**
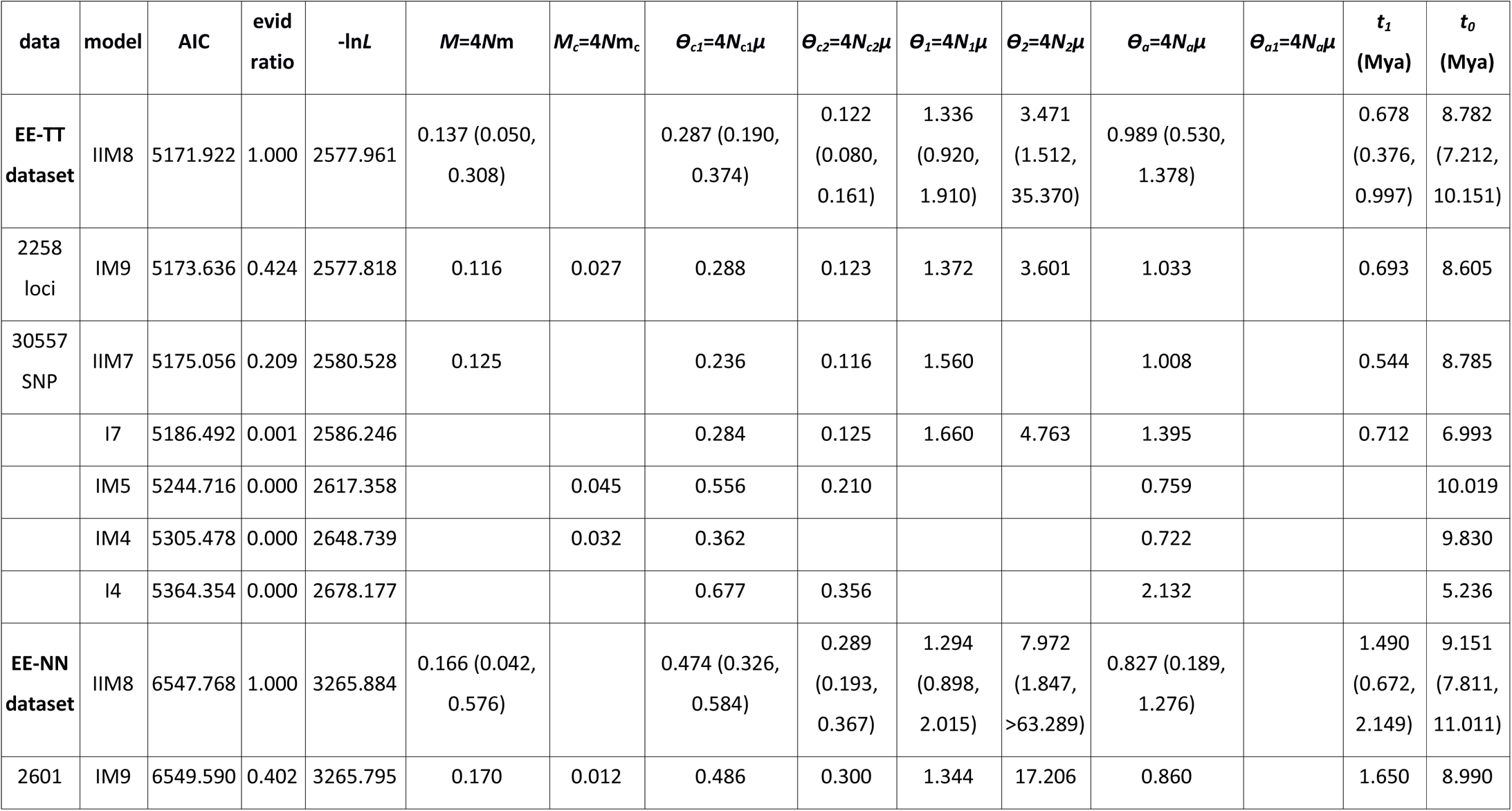
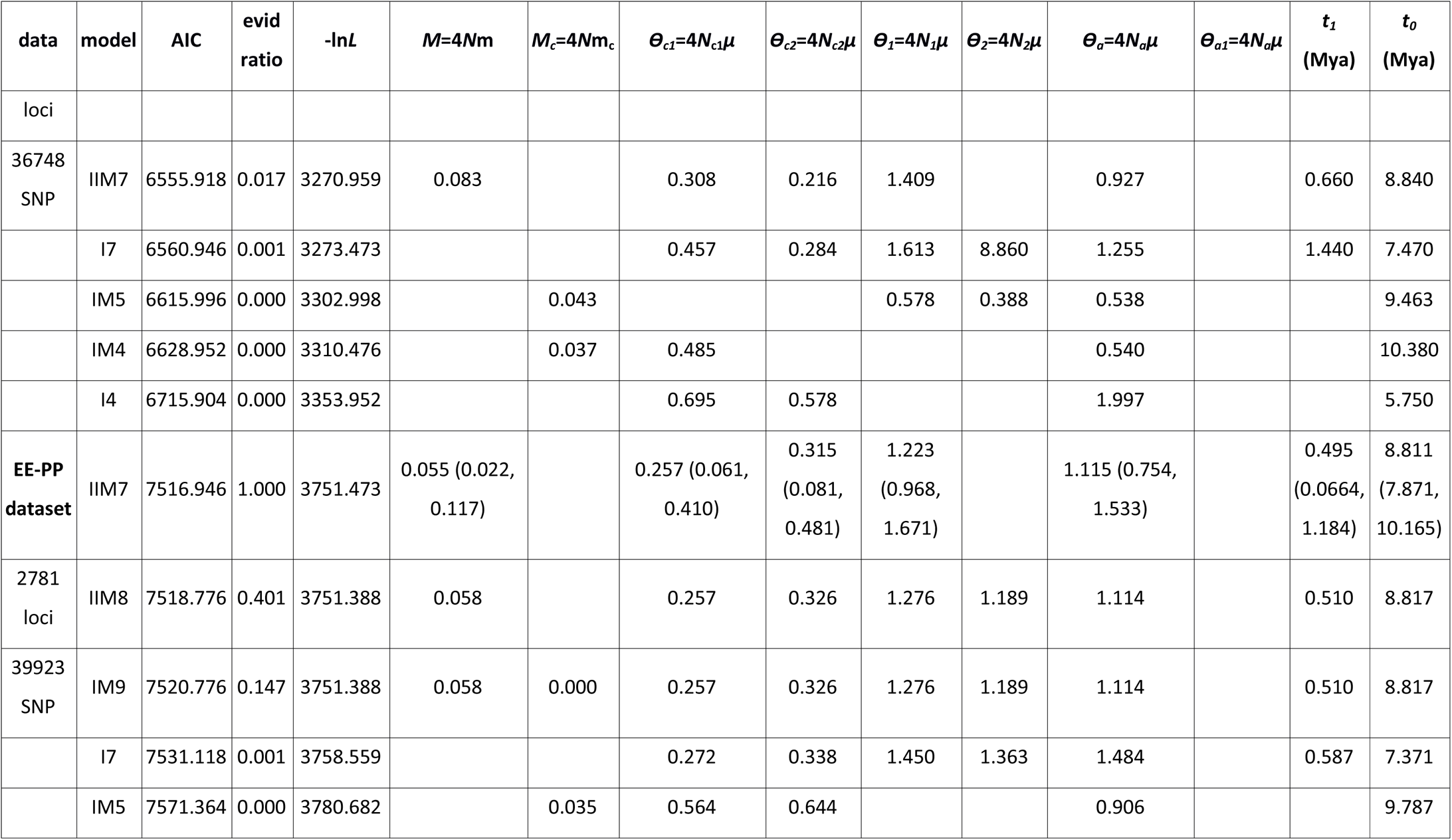
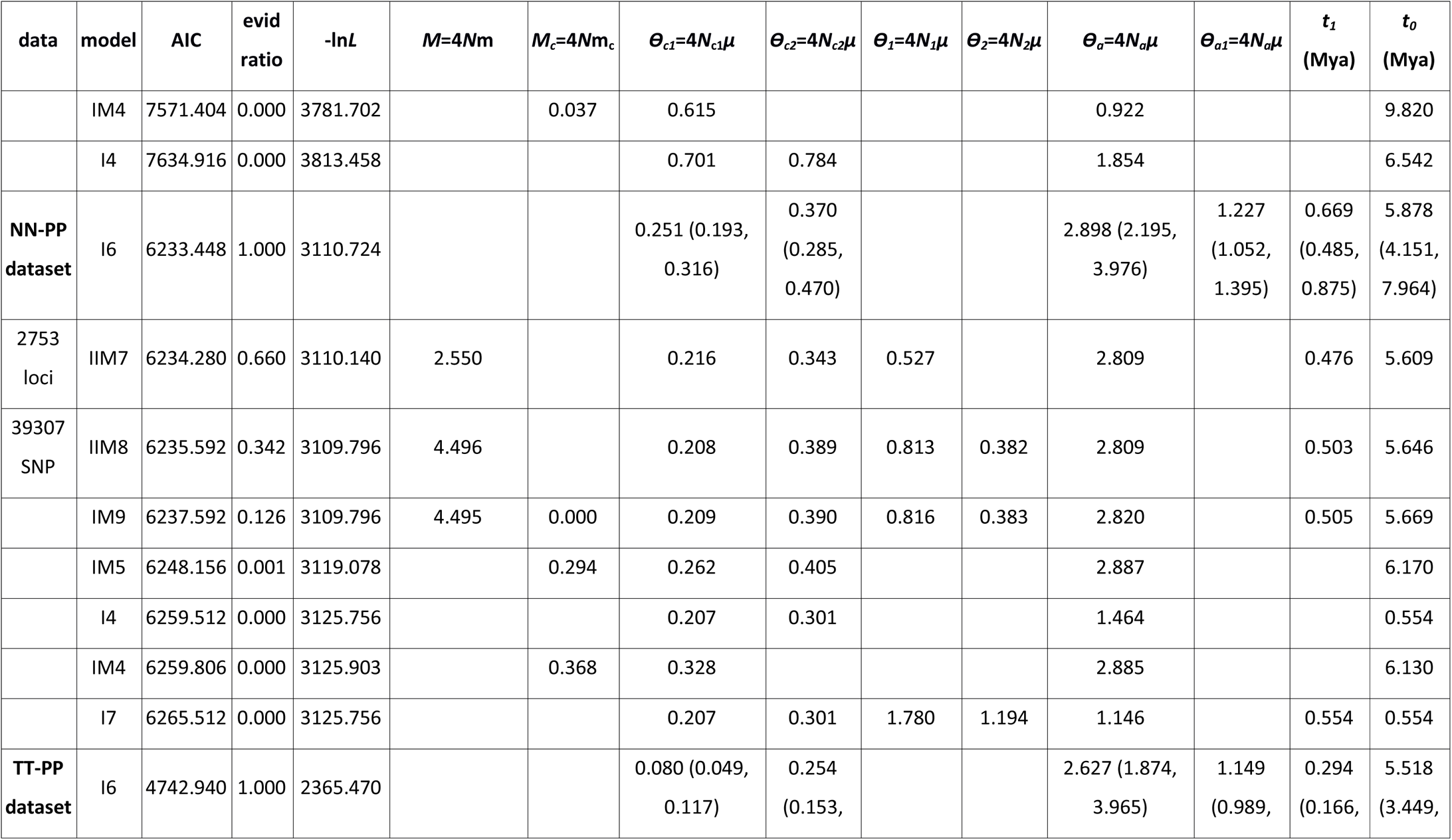
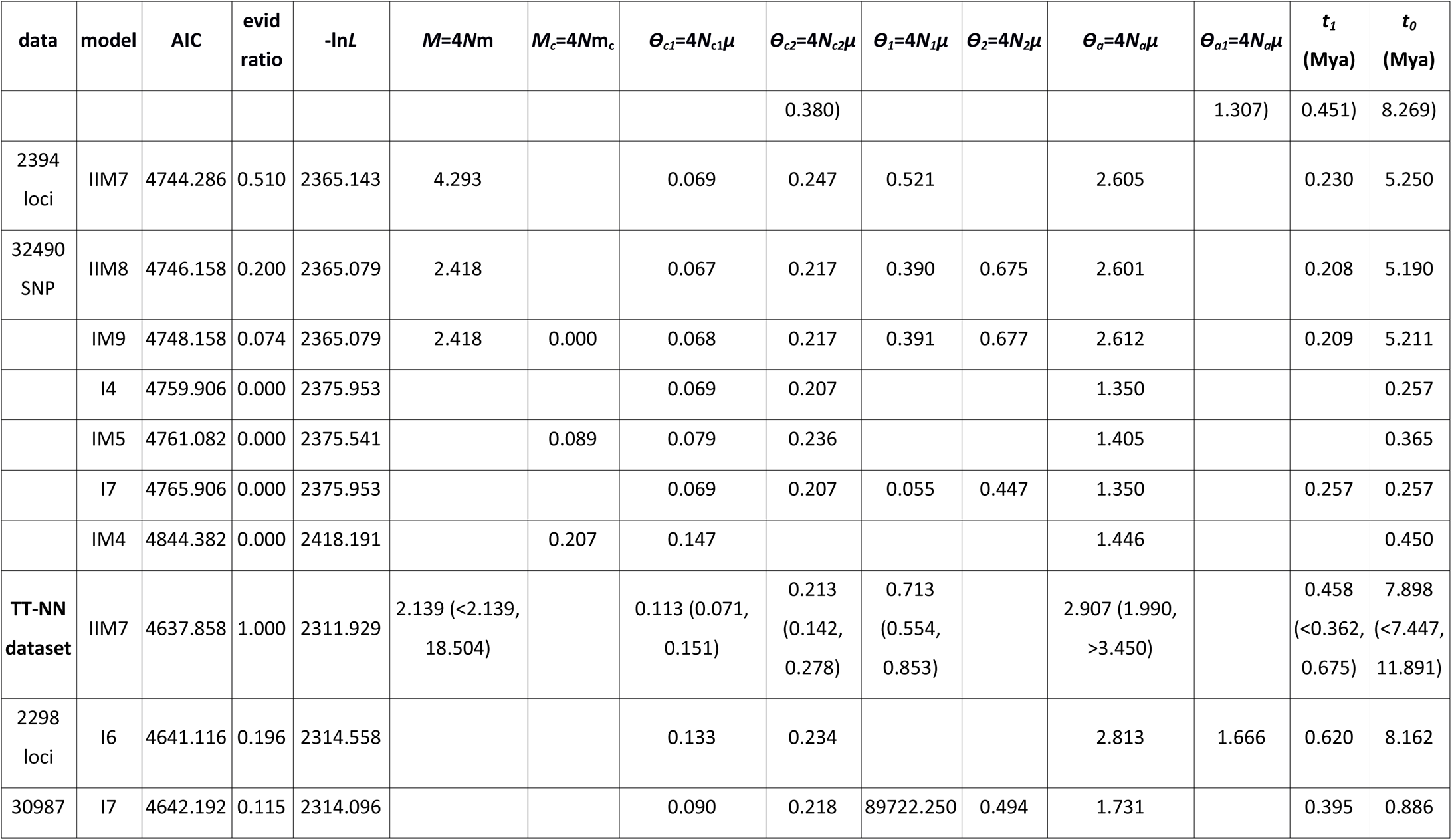
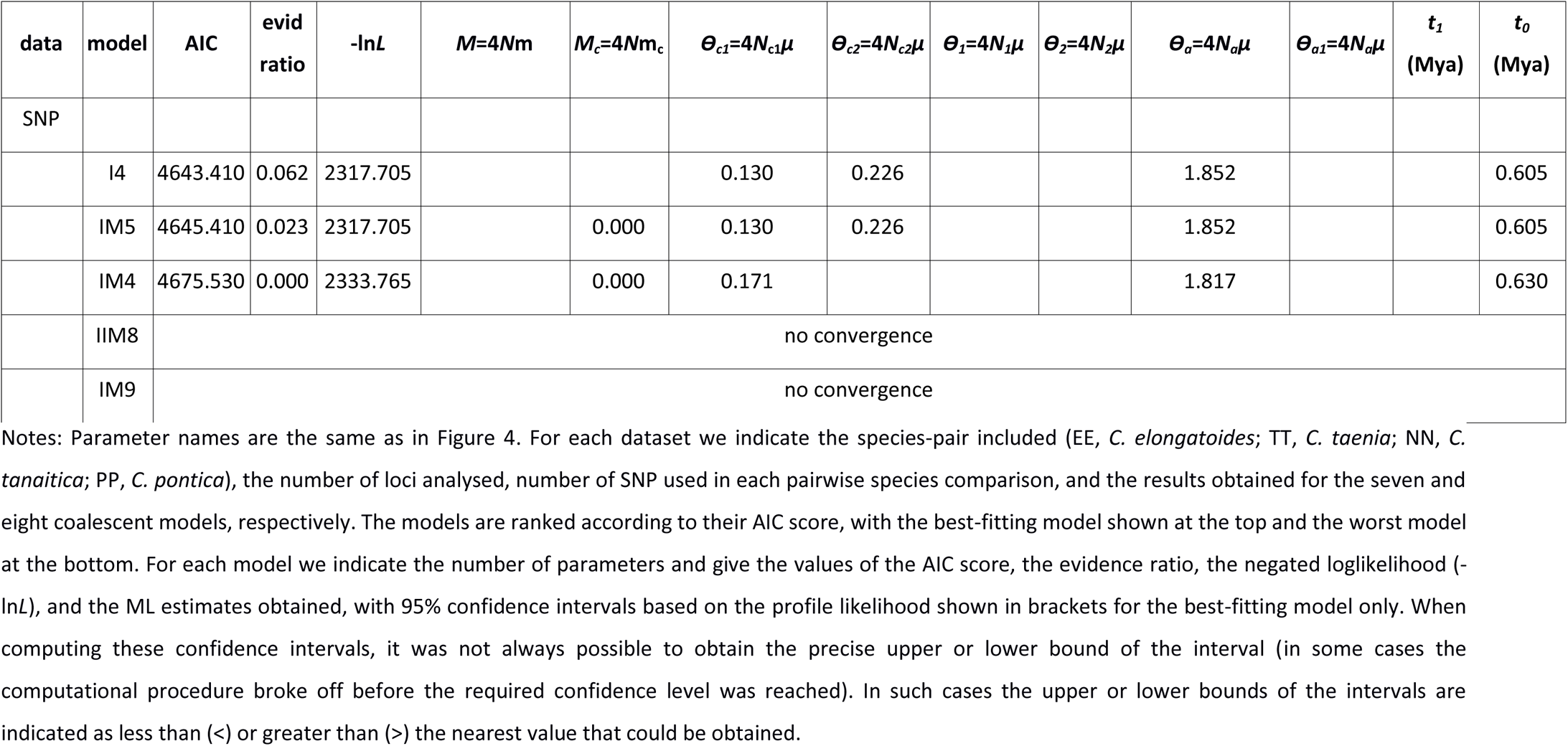
Summary of the coalescent models.

| data | model | AIC | evid ratio | -lnL | $M=4Nm$ | $M_c=4Nm_c$ | $\theta_{c1}=4N_{c1}\mu$ | $\theta_{c2}=4N_{c2}\mu$ | $\theta_1=4N_1\mu$ | $\theta_2=4N_2\mu$ | $\theta_a=4N_a\mu$ | $\theta_{a1}=4N_{a1}\mu$ | $t_1$<br>(Mya) | $t_0$<br>(Mya) |
| --- | --- | --- | --- | --- | --- | --- | --- | --- | --- | --- | --- | --- | --- | --- |
| **EE-TT dataset** | IIM8 | 5171.922 | 1.000 | 2577.961 | 0.137 (0.050, 0.308) |  | 0.287 (0.190, 0.374) | 0.122 (0.080, 0.161) | 1.336 (0.920, 1.910) | 3.471 (1.512, 35.370) | 0.989 (0.530, 1.378) |  | 0.678 (0.376, 0.997) | 8.782 (7.212, 10.151) |
| 2258 loci | IM9 | 5173.636 | 0.424 | 2577.818 | 0.116 | 0.027 | 0.288 | 0.123 | 1.372 | 3.601 | 1.033 |  | 0.693 | 8.605 |
| 30557 SNP | IIM7 | 5175.056 | 0.209 | 2580.528 | 0.125 |  | 0.236 | 0.116 | 1.560 |  | 1.008 |  | 0.544 | 8.785 |
|  | I7 | 5186.492 | 0.001 | 2586.246 |  |  | 0.284 | 0.125 | 1.660 | 4.763 | 1.395 |  | 0.712 | 6.993 |
|  | IM5 | 5244.716 | 0.000 | 2617.358 |  | 0.045 | 0.556 | 0.210 |  |  | 0.759 |  |  | 10.019 |
|  | IM4 | 5305.478 | 0.000 | 2648.739 |  | 0.032 | 0.362 |  |  |  | 0.722 |  |  | 9.830 |
|  | I4 | 5364.354 | 0.000 | 2678.177 |  |  | 0.677 | 0.356 |  |  | 2.132 |  |  | 5.236 |
| **EE-NN dataset** | IIM8 | 6547.768 | 1.000 | 3265.884 | 0.166 (0.042, 0.576) |  | 0.474 (0.326, 0.584) | 0.289 (0.193, 0.367) | 1.294 (0.898, 2.015) | 7.972 (1.847, >63.289) | 0.827 (0.189, 1.276) |  | 1.490 (0.672, 2.149) | 9.151 (7.811, 11.011) |
| 2601 | IM9 | 6549.590 | 0.402 | 3265.795 | 0.170 | 0.012 | 0.486 | 0.300 | 1.344 | 17.206 | 0.860 |  | 1.650 | 8.990 |

| data | model | AIC | evid<br>ratio | -lnL | $M=4Nm$ | $M_c=4Nm_c$ | $\theta_{c1}=4N_{c1}\mu$ | $\theta_{c2}=4N_{c2}\mu$ | $\theta_1=4N_1\mu$ | $\theta_2=4N_2\mu$ | $\theta_a=4N_a\mu$ | $\theta_{a1}=4N_{a1}\mu$ | $t_1$<br>(Mya) | $t_0$<br>(Mya) |
| --- | --- | --- | --- | --- | --- | --- | --- | --- | --- | --- | --- | --- | --- | --- |
| loci |  |  |  |  |  |  |  |  |  |  |  |  |  |  |
| 36748<br>SNP | IIM7 | 6555.918 | 0.017 | 3270.959 | 0.083 |  | 0.308 | 0.216 | 1.409 |  | 0.927 |  | 0.660 | 8.840 |
|  | I7 | 6560.946 | 0.001 | 3273.473 |  |  | 0.457 | 0.284 | 1.613 | 8.860 | 1.255 |  | 1.440 | 7.470 |
|  | IM5 | 6615.996 | 0.000 | 3302.998 |  | 0.043 |  |  | 0.578 | 0.388 | 0.538 |  |  | 9.463 |
|  | IM4 | 6628.952 | 0.000 | 3310.476 |  | 0.037 | 0.485 |  |  |  | 0.540 |  |  | 10.380 |
|  | I4 | 6715.904 | 0.000 | 3353.952 |  |  | 0.695 | 0.578 |  |  | 1.997 |  |  | 5.750 |
| **EE-PP**<br>**dataset** | IIM7 | 7516.946 | 1.000 | 3751.473 | 0.055 (0.022,<br>0.117) |  | 0.257 (0.061,<br>0.410) | 0.315<br>(0.081,<br>0.481) | 1.223<br>(0.968,<br>1.671) |  | 1.115 (0.754,<br>1.533) |  | 0.495<br>(0.0664,<br>1.184) | 8.811<br>(7.871,<br>10.165) |
| 2781<br>loci | IIM8 | 7518.776 | 0.401 | 3751.388 | 0.058 |  | 0.257 | 0.326 | 1.276 | 1.189 | 1.114 |  | 0.510 | 8.817 |
| 39923<br>SNP | IM9 | 7520.776 | 0.147 | 3751.388 | 0.058 | 0.000 | 0.257 | 0.326 | 1.276 | 1.189 | 1.114 |  | 0.510 | 8.817 |
|  | I7 | 7531.118 | 0.001 | 3758.559 |  |  | 0.272 | 0.338 | 1.450 | 1.363 | 1.484 |  | 0.587 | 7.371 |
|  | IM5 | 7571.364 | 0.000 | 3780.682 |  | 0.035 | 0.564 | 0.644 |  |  | 0.906 |  |  | 9.787 |
|  | IM4 | 7571.404 | 0.000 | 3781.702 |  | 0.037 | 0.615 |  |  |  | 0.922 |  |  | 9.820 |
|  | I4 | 7634.916 | 0.000 | 3813.458 |  |  | 0.701 | 0.784 |  |  | 1.854 |  |  | 6.542 |
| **NN-PP**<br>**dataset** | I6 | 6233.448 | 1.000 | 3110.724 |  |  | 0.251 (0.193,<br>0.316) | 0.370<br>(0.285,<br>0.470) |  |  | 2.898 (2.195,<br>3.976) | 1.227<br>(1.052,<br>1.395) | 0.669<br>(0.485,<br>0.875) | 5.878<br>(4.151,<br>7.964) |
| 2753<br>loci | IIM7 | 6234.280 | 0.660 | 3110.140 | 2.550 |  | 0.216 | 0.343 | 0.527 |  | 2.809 |  | 0.476 | 5.609 |
| 39307<br>SNP | IIM8 | 6235.592 | 0.342 | 3109.796 | 4.496 |  | 0.208 | 0.389 | 0.813 | 0.382 | 2.809 |  | 0.503 | 5.646 |
|  | IM9 | 6237.592 | 0.126 | 3109.796 | 4.495 | 0.000 | 0.209 | 0.390 | 0.816 | 0.383 | 2.820 |  | 0.505 | 5.669 |
|  | IM5 | 6248.156 | 0.001 | 3119.078 |  | 0.294 | 0.262 | 0.405 |  |  | 2.887 |  |  | 6.170 |
|  | I4 | 6259.512 | 0.000 | 3125.756 |  |  | 0.207 | 0.301 |  |  | 1.464 |  |  | 0.554 |
|  | IM4 | 6259.806 | 0.000 | 3125.903 |  | 0.368 | 0.328 |  |  |  | 2.885 |  |  | 6.130 |
|  | I7 | 6265.512 | 0.000 | 3125.756 |  |  | 0.207 | 0.301 | 1.780 | 1.194 | 1.146 |  | 0.554 | 0.554 |
| **TT-PP**<br>**dataset** | I6 | 4742.940 | 1.000 | 2365.470 |  |  | 0.080 (0.049,<br>0.117) | 0.254<br>(0.153, |  |  | 2.627 (1.874,<br>3.965) | 1.149<br>(0.989, | 0.294<br>(0.166, | 5.518<br>(3.449, |
|  |  |  |  |  |  |  |  | 0.380) |  |  |  | 1.307) | 0.451) | 8.269) |
| 2394<br>loci | IIM7 | 4744.286 | 0.510 | 2365.143 | 4.293 |  | 0.069 | 0.247 | 0.521 |  | 2.605 |  | 0.230 | 5.250 |
| 32490<br>SNP | IIM8 | 4746.158 | 0.200 | 2365.079 | 2.418 |  | 0.067 | 0.217 | 0.390 | 0.675 | 2.601 |  | 0.208 | 5.190 |
|  | IM9 | 4748.158 | 0.074 | 2365.079 | 2.418 | 0.000 | 0.068 | 0.217 | 0.391 | 0.677 | 2.612 |  | 0.209 | 5.211 |
|  | I4 | 4759.906 | 0.000 | 2375.953 |  |  | 0.069 | 0.207 |  |  | 1.350 |  |  | 0.257 |
|  | IM5 | 4761.082 | 0.000 | 2375.541 |  | 0.089 | 0.079 | 0.236 |  |  | 1.405 |  |  | 0.365 |
|  | I7 | 4765.906 | 0.000 | 2375.953 |  |  | 0.069 | 0.207 | 0.055 | 0.447 | 1.350 |  | 0.257 | 0.257 |
|  | IM4 | 4844.382 | 0.000 | 2418.191 |  | 0.207 | 0.147 |  |  |  | 1.446 |  |  | 0.450 |
| **TT-NN**<br>**dataset** | IIM7 | 4637.858 | 1.000 | 2311.929 | 2.139 (<2.139,<br>18.504) |  | 0.113 (0.071,<br>0.151) | 0.213<br>(0.142,<br>0.278) | 0.713<br>(0.554,<br>0.853) |  | 2.907 (1.990,<br>>3.450) |  | 0.458<br>(<0.362,<br>0.675) | 7.898<br>(<7.447,<br>11.891) |
| 2298<br>loci | I6 | 4641.116 | 0.196 | 2314.558 |  |  | 0.133 | 0.234 |  |  | 2.813 | 1.666 | 0.620 | 8.162 |
| 30987 | I7 | 4642.192 | 0.115 | 2314.096 |  |  | 0.090 | 0.218 | 89722.250 | 0.494 | 1.731 |  | 0.395 | 0.886 |
| SNP |  |  |  |  |  |  |  |  |  |  |  |  |  |  |
|  | I4 | 4643.410 | 0.062 | 2317.705 |  |  | 0.130 | 0.226 |  |  | 1.852 |  |  | 0.605 |
|  | IM5 | 4645.410 | 0.023 | 2317.705 |  | 0.000 | 0.130 | 0.226 |  |  | 1.852 |  |  | 0.605 |
|  | IM4 | 4675.530 | 0.000 | 2333.765 |  | 0.000 | 0.171 |  |  |  | 1.817 |  |  | 0.630 |
|  | IIM8 | no convergence |  |  |  |  |  |  |  |  |  |  |  |  |
|  | IM9 | no convergence |  |  |  |  |  |  |  |  |  |  |  |  |

Notes: Parameter names are the same as in Figure 4. For each dataset we indicate the species-pair included (EE, *C. elongatoides*; TT, *C. taenia*; NN, *C. tanaitica*; PP, *C. pontica*), the number of loci analysed, number of SNP used in each pairwise species comparison, and the results obtained for the seven and eight coalescent models, respectively. The models are ranked according to their AIC score, with the best-fitting model shown at the top and the worst model at the bottom. For each model we indicate the number of parameters and give the values of the AIC score, the evidence ratio, the negated loglikelihood (-lnL), and the ML estimates obtained, with 95% confidence intervals based on the profile likelihood shown in brackets for the best-fitting model only. When computing these confidence intervals, it was not always possible to obtain the precise upper or lower bound of the interval (in some cases the computational procedure broke off before the required confidence level was reached). In such cases the upper or lower bounds of the intervals are indicated as less than (<) or greater than (>) the nearest value that could be obtained.

Arrows along the side of model diagrams indicate the respective time periods. The population size parameter is defined as *θ*_i_= 4*N*_i_*μ*, where *N*_i_ is the effective diploid size of species i and *μ* is the mutation rate per sequence per generation, averaged over the loci included in the analysis; migration rate is defined as *M* = 4*Nm*, where *m* is the proportion of migrants per generation. Wherever an index ‘c’ accompanies the parameter name, it will always indicate the values relevant for current populations, while an index ‘a’ indicates the states of ancestral populations before the split.

Isolation-with-initial-migration model IIM7 (where the label ‘7’ indicates the number of parameters of the model) assumes that gene flow occurred from the time *t_0_* when the ancestral species split until time *t_1_* when the descendant species became completely isolated from each other. This model fitted all pairwise species datasets significantly better than both other models of Wilkinson-Herbots [36, 37], namely the isolation model I4 assuming no gene flow since the initial split of the species, and the isolation-with-migration model IM4 assuming ongoing hybridization from the initial split until the present (Likelihood Ratio Test; LRT *p* << 10^-10^ for all comparisons involving *C. elongatoides*, and *p* < 0.01 for all other species comparisons). To test if the better fit of IIM7 is due to the correct assessment that interspecific gene flow ceased recently, or merely due to allowing an additional size change at time *t_1_*, we employed five more models that allow an additional population size change and also relax the assumption of equal population sizes during the migration stage (Fig 4; [39, 40]).

Pairwise datasets of the three closely related *C. taenia, C. tanaitica* and *C. pontica* were better fitted by the isolation-with-initial-migration models (IIM7, IIM8) compared to the I4 and I7 isolation models (LRT *p* < 0.01 for all comparisons). The generalized isolation-with-migration GIM9 model gave estimates of 0 for the current migration rate and hence reduced to the IIM8 model (where these models could be fitted). However, these models consistently suggested that the analysed species were interconnected by a very intensive gene flow at a level close to one migrant sequence per generation or even more until a time *t_1_* of approximately 0.2 – 0.5 Mya. Such high levels of gene flow are considered close to panmixia (e.g. [41]) suggesting that those three taxa might have formed a single substructured species between times *t_0_* and *t_1_* and speciated only recently. To explore this possibility, we additionally fitted the isolation I6 model allowing one additional size change of the ancestral population before the isolation phase. The I6 model also estimated the speciation time at around 0.5 Mya and indeed provided a very good fit for all three datasets, being selected as the best model for two of these. Thus, coalescent analysis suggests that *C. taenia, C. tanaitica* and *C. pontica* are currently isolated and possibly speciated only recently.

The datasets of species pairs including *C. elongatoides* and any one of *C. taenia, C. tanaitica* or *C. pontica* were fitted better by the isolation-with-initial-migration model IIM8 than by isolation models I7 and I4 (LRT *p* < 0.0002 in all cases) and both isolation-with-migration models, IM4 and IM5 (LRT *p* << 10^-10^ in all cases). The IIM8 model also fitted the data better that the IIM7 model (which assumes equal population sizes during the migration stage) in the case of *C. elongatoides – C. taenia* and *C. elongatoides – C.tanaitica* species pairs (lower AIC scores and LRT *p* < 0.023 and 0.001, respectively) but not for *C. elongatoides – C. pontica* species pairs where IIM7 had a better AIC score. The divergence time estimates were consistent across species comparisons and place the initial split of *C. elongatoides* from the other species at roughly 9 Mya. It is estimated that *C. elongatoides* exchanged genes with the other species at average rates of less than 0.1 migrants per generation until time *t_1_*, for which ML estimates vary between 1.65 and 0.49 Mya depending on the data set and model but have generally quite large confidence intervals (Table 3). Given that the speciation times of the other three species are much more recent (see above) than their split from *C. elongatoides*, our results suggest that the detected gene flow occurred predominantly between *C. elongatoides* and the common ancestor of the other three species. The most complex GIM9 model did not improve the fit to the data compared to the IIM8 model (as indicated by the AIC scores of the models and by LRT; *p* > 0.59 for all cases). Moreover, the GIM9 model estimated that recent migration rates were much lower than historical ones, thus virtually converging to the IIM8 model and confirming that *C. elongatoides* has been historically exchanging genes with the other species but became isolated in recent times.

### Comparative analysis of genetic divergence between hybridizing fish species and types of reproductive isolation including hybrid asexuality

Finally, we tested the generality of our findings, implying that the hybrids’ asexuality may represent an intermediate stage of species diversification process. To do so, we investigated the general correlation between genetic divergence of hybridizing pairs of fish species and the dysfunction of their F1 fish hybrids. Dysfunction was measured in terms of infertility, inviability or asexual reproduction of hybrids (see Methods section for details how we amended the dataset from Russell’s [5] comparative study to incorporate asexuality). We found that the Kimura 2-parameter (K2P) corrected divergences of cytochrome *b* gene sequences from fish species pairs that produce asexual hybrids range from 0.051 to 0.166 (mean = 0.122; s.e. = 0.038). This appears as intermediate between those species pairs producing both sexes of hybrids fertile and viable (Russell’s [5] hybrid class 0; mean = 0.079; s.e. = 0.054) and those pairs that produce viable but infertile hybrids of both sexes (Russell’s hybrid class 2; mean = 0.179; s.e. = 0.025) (Fig 5 and Appendix S5). Although some species of the *Hexagrammos* and *Poecilia* genera were involved in more types of hybridization producing asexuals, we treated each species only once and repeated the analysis several times to account for all combinations. The Shapiro-Wilk test did not reject normality in any of the hybrid classes tested (*p* > 0.05)and regardless of species pairs considered, parental divergences in the asexual hybrid class were always significantly lower than those of Russell’s [5] class 2 (Student’s t-test, *p* < 0.01) and always significantly higher than that of Russell’s [5] hybrid class 0 (t-test, *p* < 0.05). Interestingly, the range of divergences among asexual hybrids is notably similar to the divergences of hybrids where the functionality of one sex is lower than that of the other (hybrid classes 0.5 – 1.5; mean = 0.118; s.e. = 0.040).

**Figure 5.**
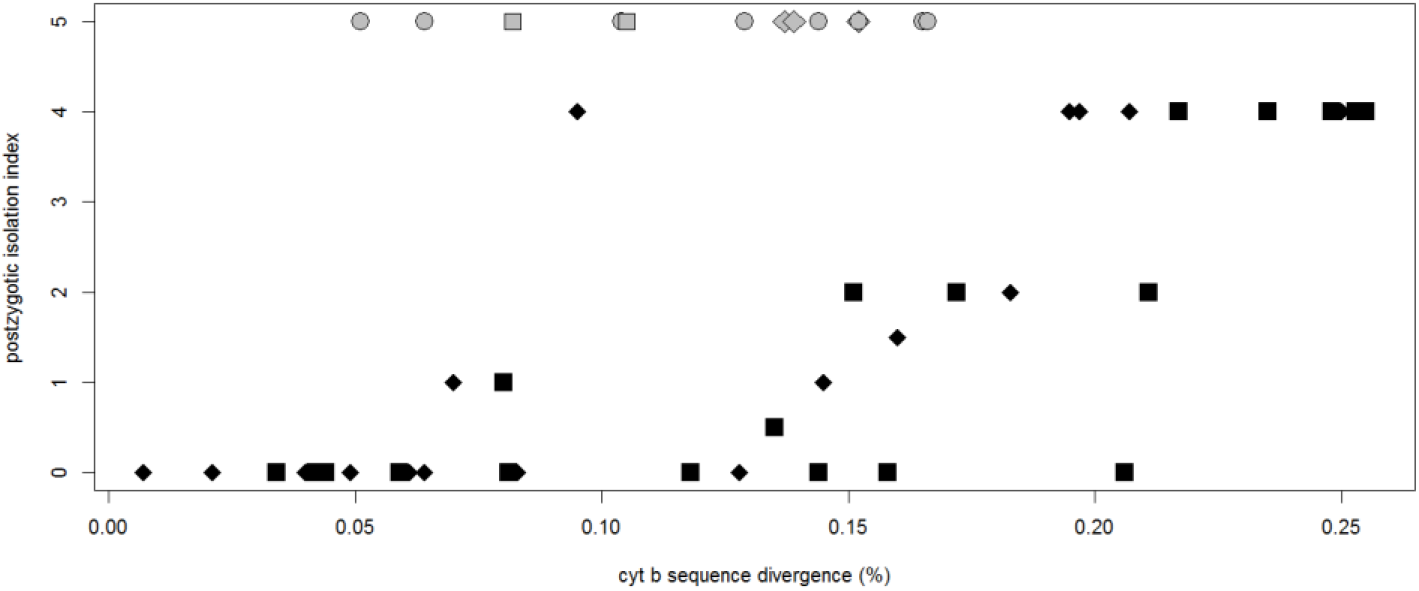
Correlation between intrinsic post-zygotic reproductive isolation and K2P corrected distances in cytochrome *b* gene between hybridizing species.

Reproductive isolation index is defined according to Russell’s study as follows: 0, both hybrid sexes are fertile; 0.5, one sex fertile, the other sometimes infertile; 1, one sex fertile, the other infertile but viable; 1.5, one sex infertile but viable, the other sometimes still fertile; 2, both sexes viable but infertile; 2.5, one sex viable but infertile, the other sex only sometimes viable; 3, one sex viable, the other missing; 3.5, one sex sometimes viable, the other not; 4, both sexes inviable; 5, hybrids of at least one sex are known to form asexual lineages (highlighted in grey color). Species pairs where one species occurred more than once in the analysis are indicated by grey diamonds *(Poeciliopsis monacha)* and grey squares *(Hexagrammos octogramus)*, respectively.

## Discussion

Natural asexual organisms are considered as treasured evolutionary models because they could provide clues about the paradox of sex. Our study offers a conceptually novel view on this phenomenon and suggests that hybrid asexuality may form an inherent stage of the speciation process and constitute an effective reproductive barrier, which arises earlier in speciation than hybrid sterility and inviability. Our conclusion is based on the following findings: (1) We demonstrated that both the intensity of interspecific gene flow as well as the likelihood of formation of asexual hybrids change with increasing genetic distance between parental species. (2) We confirmed that hybrid asexuality creates an effective reproductive barrier to interspecific gene flow. (3) We demonstrated the general applicability of our hypothesis by finding that hybrid asexuality in fish arises at lower genetic distances than complete hybrid sterility and hybrid inviability and might thus represent a primary reproductive barrier in many taxa.

### Accomplishment of speciation in spite of ongoing hybridization

The *Cobitis* species considered readily hybridize both in natural and experimental conditions [24, 28, 30], suggesting the absence of strong prezygotic RIMs. However, two independent lines of evidence suggest that contemporary interspecific gene flow between the species with overlapping ranges, i.e., *C. elongatoides* on one side and *C. taenia* and *C. tanaitica* on the other side, is unlikely and that speciation has been completed. First, according to experimental crossings of *C. elongatoides-taenia* [24, 29] and *C. elongatoides-tanaitica* (present study), hybrid females do not produce reduced gametes through the “standard” sexual process and do not produce recombinant progeny. Allozyme analyses of oocytes further argue against the production of reduced gametes through hybridogenesis; contrary to evidence from hybridogenetic vertebrates, which exclude one parental species’ genome [32–34], we found that oocytes of studied *Cobitis* hybrid females expressed allozyme alleles of both parental species. Unreduced gametes either develop clonally or occasionally incorporate the sperm’s genome leading to polyploidy, but are unlikely to enable interspecific gene flow. Hybrid males also do not appear to mediate gene flow since *C. elongatoides-taenia* hybrid males are infertile [24] and *C. elongatoides-tanaitica* hybrid males have not been observed in nature.

Second, the completion of speciation is also supported by analyses of *C. elongatoides – C. taenia* and *C. elongatoides – C. tanaitica* hybrid zones. The contact zone between *C. elongatoides* and C. *tanaitica*, covering most of the Danube watershed below the Iron Gates Gorge, is larger than the Central European *C. elongatoides – taenia* hybrid zone [30] and also involves the presence of a third, phylogenetically distant, species *C. strumicae*, which invaded Danubian stretches during postglacial expansion from the Mediterranean Basin [26]. Nevertheless, both zones have similarities with respect to major evolutionary aspects. Polyploid asexuals dominate in both zones (in the Danube drainage, they also invaded rivers inhabited by *C. strumicae* [42] and diploids within both zones could consistently be assigned into two types. The first type was represented by one or the other pure sexual species, the second type was represented by hybrids. Unlike classical patterns where the distribution of the hybrids’ values of the *q* parameter follow a continuum, the *q* index for *Cobitis* was sharply unimodal around intermediate values and most hybrids were unambiguously assigned as F1 (NewHybrids software, *p* > 0.95). Only in three Danubian diploid lineages we could not reject the B1 state although the F1 state was still preferred. Interestingly, these three lineages belong to the old clonal Hybrid clade I and possess a number of private microsatellite alleles, suggesting that their assignment by both the Structure and NewHybrids software might have been affected by mutational divergence from contemporary sexual species (see e.g. [43]). Most diploid hybrids from both zones also clustered into lineages of (nearly) identical individuals that may be assigned as clones and all but one (MLG B) diploid hybrid lineages belong to an ancient asexual Hybrid clade I arisen about 0.35 Mya. Contemporary hybrids of *C. elongatoides* thus do not appear able to produce recombinant progeny mediating interspecific gene flow. As *Cobitis* species do not notably differ in ecological requirements, thereby excluding the possibility of strong extrinsic postzygotic isolation, we suggest that the asexuality of hybrid females together with the sterility of hybrid males is the primary reproductive barrier separating these species.

Although we found no evidence of contemporary gene flow between *C. elongatoides* on one side and *C. taenia* and *C. tanaitica* on the other side, coalescent analysis of RNAseq data suggested that hybrids that were able to mediate gene flow must have existed in the past. We clearly rejected isolation models, suggesting that gene flow existed after the initial divergence of *C. elongatoides* from *C. taenia*, *C. pontica* and *C. tanaitica*. Interestingly, datasets including *C. elongatoides* were significantly better fitted by isolation-with-initial-migration models than by the simpler isolation-with-migration models, which implies that gene flow involving *C. elongatoides* occurred in the historical period after its initial divergence around 9 Mya but did not occur after *t_1_*, suggesting the isolation of contemporary populations (Fig 4 and Table 3). As the applied coalescent models assumed gene flow since the initial split of *C. elongatoides* but compared only pairs of extant species, it is reasonable to conclude that the inferred gene flow occurred between *C. elongatoides* and the common *C. taenia – C. tanaitica – C. pontica* ancestor that diversified only recently around ca 0.5 Mya. Unfortunately, the large confidence intervals of the *t_1_* estimates prevent an unambiguous conclusion as to whether nuclear gene flow continued after the *C. taenia – C. tanaitica – C. pontica* speciation. Conversely, we found no evidence of gene flow between the closely related *C. taenia*, *C. tanaitica* and *C. pontica*, where coalescent models suggested their recent Speciation, probably during the last 0.5 Mya

The present results are generally consistent with our previous study based on nine nuclear and one mtDNA loci but two differences were noted. First, Choleva et al. [31] found significant traces of mitochondrial but not nuclear *C. elongatoides – C. tanaitica* gene flow, which led to the somewhat paradoxical impression that the nucleus has not been affected by hybridization while *C. tanaitica’s* mitochondrion has suffered one of the most massive introgressions among animals (a complete mtDNA replacement). The current analysis of the extended dataset indicated gene flow in the nuclear compartment too, which is more plausible biologically. Second, the previous model-based inference indicated *C. taenia – C. tanaitica* gene flow in nucleus, which was at odds with the available field data because no *C. taenia - tanaitica* hybrids have ever been observed in nature and the ranges of both species are closely adjacent but sympatry has not been documented [28]. The present analysis is therefore more in line with current knowledge about *Cobitis* and the discrepancy with the previous analysis of only few loci may potentially reflect the tendency of Bayesian IM algorithms to inflate estimates of gene flow when the number of loci is low and splitting times are recent [44, 45].

Of course, any model-based inferences should be treated with caution. For example, having analysed only pairs of species, our models may be affected by intractable interactions with other species. IM models are robust to this type of violation, for low or moderate levels of gene flow [46], and the inferred lack of hybridization between *C. taenia – C. tanaitica – C. pontica* implies that the pairwise comparison of any one of these species with *C. elongatoides* is unlikely to be affected by the existence of the other species. However, other problems may be important, such as uncertainty regarding the relative mutation rates of the different loci (which have been estimated by comparing them with the outgroup), additional population size changes, geographical structure or varying rather than constant intensity of gene flow between *t_0_* and *t_1_*. Furthermore, coalescent models assume evolution under neutrality or negative selection, which effectively decreases the locus-specific mutation rate [47], but other types of selection might have affected our data. It is therefore a great advantage that model-based inferences are supported by independent types of data. The existence of gene flow between *C. elongatoides* and other species is supported by fixation of an C. *elongatoides-like* mitochondrial lineage in *C. tanaitica* [31]. On the other hand, small but non-zero divergences between contemporary *C. elongatoides* and *C. tanaitica* mtDNA haplotypes, clonal reproduction of the hybrids studied, and the apparent lack of introgressive hybridization in hybrid zones imply that such gene flow ceased and does not occur at present.

### Simultaneous evolution of asexuality and RIMs

Several complementary approaches showed that the initial divergence of *C. elongatoides* was coupled with the production of hybrids whose reproductive mode enabled more or less intensive gene flow with other three species or common ancestor these species. However, as *C. elongatoides* diverged from these species, the introgressions became restricted since the major type of hybrids became asexual, which may not effectively mediate the gene flow. Such patterns conform to the “balance hypothesis” [22], which predicts that hybridization between gradually diverging species would initially produce mostly sexual hybrids while successful asexuals would arise at intermediate stages when a hybrid’s meiosis is disrupted but fertility is not yet significantly reduced. Simultaneously, this scenario is consistent with the gradual decline in the species’ capability of introgressive hybridization that is expected to evolve along the speciation continuum from weakly separated entities towards strongly isolated species (e.g. [1]). The establishment of asexual hybrids and the gradual accumulation of intrinsic postzygotic RIMs thus appear to be interconnected processes that correlate with divergence of hybridizing species.

Crossing experiments support this idea since distantly related *C. elongatoides* and *C. taenia* produced only clonally reproducing F1 females and infertile F1 males [24], while mating of the closely related *C. taenia* and *C. pontica* species (this study) resulted in fertile F1 progeny of both sexes that mostly produced recombinant progeny. Such data also conform to the empirical observation that one sex often acquires infertility earlier than the other one [5, 9]. Unfortunately, lack of data on *Cobitis* sex chromosomes prevent speculations as to whether infertility of hybrid males could be related to Haldane’s rule or other processes such as the increased sensitivity of spermatogenesis compared to oogenesis to perturbations in hybrids [48]. Interestingly, we noted that some *C. taenia- pontica* female hybrids reproduced clonally. This observation extends the list of previously published asexual lineages and suggests that the emergence of asexual hybrids is a taxonomically widespread phenomenon within the entire family *Cobitidae*, involving several independent species pairs. These include *C. taenia-pontica* (laboratory hybrids from this study), *C. elongatoides-taenia* and *C. elongatoides-tanaitica* [28] as well as two cases from other related genera of the family [49, 50]. Such a widespread and independent emergence of asexuality is not consistent with the phylogenetic constraint hypothesis [51, 52], which assumes that the initiation of hybrid asexuals may result from predisposition of only certain parental species or populations to produce such asexuals. Instead, our data are consistent with the Balance Hypothesis and further indicate that hybrid asexuality constitutes an intermediate stage of speciation continuum that may be viewed as a special case of Dobzhansky-Muller incompatibilities. In this respect, it is noteworthy that hybrid asexuality is commonly observed only in one direction of cross [53, 54], which is similar to classical patterns observed in hybrid infertility and inviability.

Performed comparative analysis further suggests that the scenario revealed in spined loaches can be extrapolated to other taxa. The inclusion of known asexual hybrids into Russell’s [5] study of fish hybrids showed that asexual hybrids appear at intermediate levels of parental divergences between species pairs producing fertile and viable hybrids and those producing infertile hybrids of both sexes (Fig 5, Appendix S5). In fact, the divergence of asexual-producing species pairs is similar to those producing hybrids with lowered fertility of hybrids of one sex (Russell’s hybrid classes 0.5 - 1.5), which is in line with the observation that populations of asexual fish often consist of females only. While Russell [5] corroborated that the severity of a hybrid’s dysfunction correlates with the divergence of the hybridizing fish, our amendment suggest that hybrids with a skewed reproductive mode towards asexuality tend to appear at an intermediate stage of the diversification process. The relatively wide range of divergence for the asexual hybrids also agrees with the expectation that accumulation of RIMs follows a variable-rate clock and that different types of incompatibilities accumulate in a noisy manner [11]. Similarly in reptiles, parental species known to produce asexual hybrids typically appear to be somewhat genetically distant rather than being each other’s closest relatives [55–57], and, divergences among hybridizing species that produce asexual hybrids generally appeared significantly higher than among those producing bisexual hybrids [58].

Interestingly, there also appears to be a continuum in the ability of species pairs to produce hybrid asexuals. At one end, there are dynamically hybridizing species pairs that produce diverse assemblages of asexual hybrids, e.g. *Cobitis* [24], *Pelophylax* [51], *Poeciliopsis* [19]; whereas at the other end there are monoclonal hybrid lineages stemming from one or a few ancient events while attempts to cross their contemporary sexual relatives do not produce clones, e.g., *Poecilia* [59]. *Phoxinus* may represent an intermediate case, where historical hybridizations produced a highly diverse asexual assemblage but new clones can no longer be produced [60]. Similarly, hybridization of some species pairs has been documented to produce only asexual hybrids (e.g. *Cobitis elongatoides-taenia)* while clonal and sexual hybrid females co-occur and fertile hybrid males may exist in other hybridizing pairs (e.g. *Fundulus, Rutilus rutilus x Abramis bramma)*. We suggest that such a variety of patterns may reflect that different species pairs proceeded different distances along the same continuum.

To conclude, we found that the speciation continuum may contain an inherent, previously unnoticed, stage of asexuality. The production of asexual rather than sexual hybrids helps to establish an effective barrier to gene flow even in the absence of other typical forms of RIMs. Asexuality is phylogenetically widespread but rare phenomenon in animals. However, given that asexual lineages typically have a short life span [61] and that the phase when diverging species could produce asexual hybrids is transient, it is possible that many currently reproductively incompatible species might have historically produced asexual hybrids that have gone extinct. The stage of hybrid asexuality thus certainly requires close attention in research since it might have represented an important mechanism in the speciation of many groups capable of creating asexuals, including arthropods, vertebrates or plants.

## Materials and Methods

All experimental procedures involving fish were approved by the Institutional Animal Care and Use Committee of the Institute of Animal Physiology and Genetics AS CR (IAPG AS CR), according with directives from State Veterinary Administration of the Czech Republic, permit number 124/2009, and by the permit number CZ 00221 issued by Ministry of Agriculture of the Czech Republic. Crossing experiments were conducted under the supervision of L. Choleva, holder of the Certificate of competency according to §17 of the Czech Republic Act No. 246/1992 coll. on the Protection of Animals against Cruelty (Registration number CZ 02361), provided by Ministry of Agriculture of the Czech Republic, which authorizes animal experiments in the Czech Republic.

1. Material: Fig 1 and Table 2 indicate the origin of investigated fish specimens used in this study. A priori classification of captured specimens into taxonomic units (species and hybrid types) was based on previously verified diagnostic allozyme loci and PCR-RFLP method or sequencing of the first intron of the nuclear S7 gene following [28]. Ploidy was routinely estimated from gene dose effect using allozymes as described in [28] and also by flow cytometry in cases where we disposed sufficient amount of fixed tissue.

2. Detection of current introgression in *C. elongatoides – C. tanaitica* hybrid zone.

818 individuals were genotyped and those identified as diploids were subsequently subject to analysis of nine microsatellite following the protocols of [24, 62]; multiplex 1: loci Cota_006, Cota_010, Cota_027, Cota_068, Cota_093, Cota_111; multiplex 2: loci Cota_032, Cota_033, Cota_041. We also amplified and sequenced 1190 bp fragment of the cytochrome *b* gene according to [28] in a subset of diploid individuals. The GenAlEx 6.5 software [63] was used to identify clusters of individuals sharing the same microsatellite multilocus genotype (MLG) and to calculate the probability, conditional on observed variability studied loci, that individuals sharing the same MLG arose by independent sexual events (see [30] for details). While some MLG may represent distinct clones, others may belong to the same clonal lineage (the so called ‘multilocus lineage’ (MLL) and differ only by scoring errors or post formational mutations. We used the approach described in [30] to identify groups of MLGs forming such mutilocus lineages. Phylogenetic relationships among newly discovered and previously published mtDNA haplotypes [30, 42] were reconstructed with the median-joining network [64], and drawn using NETWORK software (http://www.fluxusengineering.com/netwinfo.htm).

2.2 Detection of interspecific gene flow: microsatellite and allozyme data were used to detect admixed individuals from the *C. elongatoides – C. tanaitica* hybrid zone by two Bayesian clustering methods that are based on different algorithms but have similar power for hybrid detection [65]. We first used admixture model implemented in Structure 2.3.3 [66] to compute the parameter q, i.e. the proportion of an individual’s genome originating in one of the two inferred clusters, corresponding to the parental species *C. elongatoides* and C. *tanaitica*. The analysis was based on runs with 10^6^ iterations, following a burn-in period of 5*10^4^ iterations. Ten independent runs for number of populations varying from *K*= 1 to *K*= 10. The best value of *K* was chosen following the method proposed by Evanno et al. [67], with Structure Harvester [68], which takes into account the rate of change in the log likelihood between successive *K* values.

Second, the Bayesian clustering method implemented in NewHybrids 1.1 [69] was used to compute the posterior probability that an individual in a sample belongs to one of the defined hybrid classes or parental species. The eight genotypic classes investigated were: *C. elongatoides, C. tanaitica*, F1 hybrid, F2 hybrid, and two types of backcross to either *C. elongatoides* or *C. tanaitica*. The two backcross types included those having 75 % of their genome originated from the backcrossing species (B1 generation) and those having 95 % of their genome originated from the backcrossing species (further-generation backcrosses). Posterior distributions were evaluated by running five independent analyses to confirm convergence. We started with different random seeds, performed 10^4^ burn-in iterations and followed by 500,000 Monte Carlo Markov Chain iterations without using prior allele frequency information. Analyses were run for four combinations of prior distributions (uniform or Jeffreys for *θ* and *π* parameters) to explore the robustness of the results [69].

To minimize the effect of clonal propagation, we used only one randomly chosen representative of each unique MLL and repeated the analysis several times to check its robustness against the particular choice of MLL representatives [30]. We also repeated those analyses with either Cota_006 or Cota_041 locus removed due to their possible linkage [62]. Locus Cota_027 was removed since it does not amplify in *C. elongatoides*.

3. Experimental analysis of reproductive modes of hybrids: We performed crossing experiments under conditions described in [24] and genotyped the parents and their progeny using the microsatellite multiplex 1 to verify their mode or heredity. Two types of crossing were performed:

3.1 First, we analyzed the reproductive mode of naturally occurring diploid and triploid hybrids by individually crossing wild caught EN and EEN females (see Table 1) with males of the sexual species. Several authors [32–34] also reported more complex reproductive modes when one parental genome is excluded prior to meiosis while the other is clonally transmitted. This type of so-called hybridogenetic reproduction leads to reduced, albeit non-segregating gametes and may be detected by comparison of allozyme profiles of somatic and germinal tissues. In order to test for premeiotic genome exclusion in *Cobitis* hybrids, we have therefore compared allozyme profiles of somatic tissues of hybrid females with those from their ova using the standard allozyme methods adapted for *Cobitis* [28].

3.2 The second type of experiment represents a part of long-term project aimed at analyses of *de novo* created hybrids among different *Cobitis* species. We individually crossed males and females of different species, reared their F1 progeny until sexual maturity and combined such progeny to produce backcross or second filial generation. Between 2004 and 2009, *C. elongatoides-taenia* hybrids were successfully backcrossed and the results were published [24]. In 2006, successful spawning of several *C. pontica – C. taenia* couples took place. After reaching the maturity, *C. pontica- taenia* F1 hybrids were combined for spawning and we obtained three clutches of their F2 progeny, which were analyzed (families No. 1 and 17 in Table 1 and S1).

4. Detection of historical admixture: In order to test for periods of historical or contemporary gene flow on a genome-wide scale, we applied coalescent models to SNP data from mRNA.

4.1. mRNA sequencing and assembly: the mRNA sequencing concerned liver and oocyte tissues of 2 individuals of each C. taenia, *C. elongatoides, C. tanaitica* and *C. pontica* species as well as an outgroup species *C. strumicae* belonging to the subgenus Bicanestrinia sampled from the isolated Sredecka River (Fig 1). RNA isolations were made with the Ambion^®^ ToTALLY RNA™ Total RNA Isolation Kit. cDNA libraries were constructed by SMARTer PCR cDNA Synthesis Kit (Clontech) according to the manufacturer’s instructions with the following exceptions: modified CDS-T22 primer (5’-AAGCAGTGGTATCAACGCAGAGTTTTTGTTTTTTTCTTTTTTTTTTVN-3’) was used instead of 3’ BD SMART CDS Primer II A; first strand synthesis time was prolonged to 2 hours; and PCR conditions during second strand synthesis were modified (denaturation: 95°C/ 2 min, amplification: 18 times 95°C/ 10 sec, 65°C/ 30 sec, 72°C/ 3 min, final extension: 72°C/ 5 min). cDNA was normalized by Trimmer cDNA normalization Kit (Evrogen, Moscow, Russia). 1 μg of each normalized cDNA was used for sequencing library preparations according to the Roche Rapid Library Preparation Protocol (Roche, Welwyn Garden City, UK). Libraries were tagged using MID adapters (Roche), pooled (usually 2 samples per one large sequencing region) and sequenced using GS FLX+ chemistry (454 Life Sciences, Roche).

Initial 1886536 reads (648620753 bp) from *C. taenia* were filtered to remove low quality part of reads, adaptors and primer sequences using Trimmomatic software [70]. Technical PCR multiplicates were removed using cdhit-454 software [71]. Resulting 1707769 reads (568470258 base pairs) were used for cDNA assembly using Newbler (Software Release: 2.6 20110517_1502) with parameters: minimum read length: 40; minimum overlap length: 40; minimum match: 90 %; minimum contig length 300 bp. Assembled loci were blasted against *Cobitis takatsuensis* mitochondrion (NC_015306.1) using blastn [72] and contigs matching to mitochondrion were removed from subsequent analysis in order to analyse the phylogeny and gene flow in nucleus only. Correct detection of SNPs may be compromised by undetected paralogy, copy number variation, and repetitive parts of genomes, which tend to introduce spurious heterozygotes calling based on between-paralogue variation [73]. We removed from analysis all contigs, where read mapping was not unique. We further excluded contigs indicative of paralogous mapping, where we observed excessive heterozygosity with identical heterozygotic states being observed simultaneously at same SNP positions of distant outgroup (C. *strumicae)* as well as all scored ingroup species. First assembly (28338 cDNA contigs, 31831582 bp with N50 1301 bp and 95.39 % bases with quality score Q40 and better) was cleaned for duplicates and paralogs, which left us with transcriptome of 20385 potential mRNAs (average length 1096.5 bp, total length 22355325 bp and N50 1246 bp).

On this reference transcriptome we separately mapped reads from all species including outgroup and two individuals from each ingroup species, with the Newbler software. For each individual, all highly confident SNPs with sufficient coverage were inserted into the database using our own SQL scripts. To detect if a particular SNP was sequenced in any animal, we prepared new version of the reference transcriptome with the bases replaced by N in positions where the particular SNP was detected. This new transcriptome was then used for new mapping and detection of SNP presence/absence in all individuals. Given that the reference transcriptome was prepared from *C. taenia* but more distant species were subsequently mapped onto it, our approach introducing ‘N’ on variant positions minimized the unbalanced mapping of more distant species onto the model transcriptome. Finally, we sorted identified SNPs from each locus into individual-locus-specific matrices.

4.2 Detection of interspecific gene flow from SNP data was based on recently introduced coalescent-based maximum-likelihood methods [36–40] which are computationally inexpensive and estimate simultaneously the population sizes and migration rates as well as population splitting times, including some scenarios of time-variable migration rates. The following eight models, which are characterized in Fig 4, were fitted to the data:

a) strict isolation models (labelled I4, I6 and I7) with four, six or seven parameters, respectively, assuming that an ancestral population split into two isolated descendant populations. The I6 and I7 models allow for one additional change of size of either the ancestral or the descendant populations, respectively.

b) “isolation with migration” models (IM4 and IM5) with four or five parameters, respectively, where an ancestral population of size *θ*_a_ split at time *t_0_* into two descendant populations of equal (IM4) or unequal (IM5) sizes interconnected by gene flow at rate *M_c_;*

c) “isolation with initial migration” models (IIM7 and IIM8) with seven or eight parameters, respectively, assuming that an ancestral population of size *θ*_a_ split at time *t_0_* into two descendant populations (of equal or unequal sizes depending on the model) interconnected by gene flow at rate M, which lasted until time *t_1_* since when two descendant populations of sizes *θ*_c1_ and *θ*_c2_ evolved in isolation until the present;

The most complex “generalized isolation-with-migration” model with nine parameters (GIM9) assumes that an ancestral population of size *θ*_a_ split at time *t_0_* into two descendant populations of sizes *θ*_1_ and *θ_2_* interconnected by gene flow at rate *M* until *t*_1_, from which time onwards both species are at their current sizes *θ*_c1_ and *θ*_c2_ and gene flow occurs at its current rate *M*_c_. Note that all models described above are nested within the GIM9-model.

All models were ranked according to their AIC score and we also calculated the evidence ratio for each model, providing a relative measure of how much less likely a given model is compared to the best-fitting model, given the set of candidate models considered and the data [69]; Table 3. In addition, Likelihood Ratio tests were performed to compare pairs of nested models, where we have assumed that the use of the χ2 distribution with the appropriate number of degrees of freedom is conservative ([38, 39, 74]).

Because the currently available implementation of the above models allows the analysis of only two species at once, we prepared separate datasets, each including two of the analyzed species *C. elongatoides, C. taenia,, C. tanaitica* and *C. pontica* The datasets for each pairwise comparison were represented by locus-specific alignments of SNP positions (we did not use invariant positions) with three rows – two rows corresponded to sites with successfully resolved allelic states in both compared species and the third row contained SNPs of the outgroup species *C. strumicae*. The coalescent models assume free recombination among loci but no recombination within loci and require three sets of data for each pairwise species analysis. These consist of the number of nucleotide differences between pairs of sequences sampled a) both from species 1, b) both from species 2, and c) one from each species. To estimate all model parameters simultaneously, data from all three types of sequence pairs must be included but each pair of sequences must come from a different, independent locus [37]. Therefore, we randomly divided the analyzed loci into three non-overlapping subsets. The intraspecific data were simply calculated as the number of heterozygous positions at each locus, which in fact represents the number of nucleotide differences between a pair of alleles brought together by segregation into a sampled individual. The preparation of the third dataset was more complex since it requires the comparison of two haploid sequences from different species, while our sequences originated from diploid individuals often possessing multiple heterozygotic positions with unknown phase. Therefore, we extracted the longest possible alignment of SNPs where both compared individuals (species 1 and species 2) had at most one heterozygous position by trimming the per-locus alignments of SNPs (similar to the trimming procedure done in [75]). Subsequently we randomly phased the remaining heterozygous SNP markers into either one or the other base and compared the number of differences between both species (this procedure in fact represents the comparison of two alleles drawn from species 1 and 2 respectively while keeping the allelic states resolved in the outgroup on a shortened alignment). In order to incorporate intrapopulation variability into our estimates, we combined the data from two individuals of each species. In doing so, we randomly sampled each locus from either one or the other individual (assuming free recombination among loci) but we always kept the sequence of SNPs within each locus from a single randomly selected individual to avoid the introduction of false recombination.

The relative mutation rates at all loci were estimated by comparison with an outgroup species [76]. *C. strumicae* was used as outgroup, whose divergence time from the ingroup was set to 17.7 Mya according to the penalized likelihood time-calibrated tree of [35]. Computation of 95% confidence intervals was based on the profile likelihood as described in [39]. Because of the computational time required, such confidence intervals were provided only for the best-fitting model for each dataset. In some cases, the likelihood profiling broke down prematurely, since forcing one parameter too far from its maximum-likelihood estimate rendered the whole model impossible to fit. In such cases we indicate the nearest value that could be obtained, keeping in mind that the true 95% CI should be wider (Table 3).

5. The general applicability of patterns revealed in *Cobitis* was tested by a comparative study of published sexual and asexual fish hybrids. Russell [5] investigated the correlation between the genetic divergence between hybridizing fish species (distances in the cytochrome *b* gene corrected with the K2P model) and the level of postzygotic isolation, which was categorized by an index ranging from 0 (both hybrid sexes fertile) to 4 (both sexes inviable). The index value 2 assigned the stage when both hybrid sexes are viable but infertile, therefore preventing any effective interspecific gene flow. We amended Russel’s study by introducing additional value (5) of postzygotic isolation index to those fish hybrids that have been documented to transmit their genomes clonally (gynogenesis, androgenesis) or hemiclonally (hybridogenesis). Altogether, performed literature search lead us to 13 cases of fish asexual hybrids, which were added to the database of Russell [5]. In single case we modified Russell’s [5] data since he assigned *R. rutilus x A. bramma* hybrids with postzygotic isolation index 0.5 but Слынько [77] showed that such hybrids produce clonal gametes and can reproduce via androgenesis. Therefore, *R. rutilus x A. bramma* hybrids were assigned with index value 5. In accordance with Russell’s data, the cytochrome *b* gene divergence was calculated from available sequences of parental taxa using the K2P correction using the Mega 5.0 software [78] (Appendix S5). The genetic distances of the group 5 was compared with the other types of hybrids using the t-test after the normality of data was evaluated with the Shapiro-Wilk test.

To avoid phylogenetic dependence, Russell [5] considered each species in one cross only. Published cases of asexual hybrids concern non-overlapping species pairs with three exceptions. In *Cobitis, C. elongatoides* has been involved in at least two crosses leading to naturally occurring asexuals (C. *elongatoides-taenia* and *C. elongatoides-tanaitica)* but since the original *C. tanaitica-like* mitochondrion has been lost (see above), we considered *C. elongatoides – C. taenia* cross only. On the other hand, *Hexagrammos octogramus* and *Poeciliopsis monacha* produce asexual hybrids by mating with two *(H. otaki* and *H. agrammus)* and three *(P. lucida, P. occidentalis, P. latipina)* congenerics, respectively. Therefore, we performed the t-tests several times with only one such cross per species.

## Acknowledgements

The authors are profoundly obliged to Roger Butlin and Diego Fontaneto for their very inspiring comments during the MS writing. We also thank Sarka Pelikanova, Jana Cechova *(in memoriam)*, Koen DeGelas and Bart Hellemans for help with DNA and RNA in the laboratory, and Jana Kopecka for help with the allozyme analysis. Jan Kotusz and Petya Ivanova assisted during the field work and we are very grateful to them too.

## Author contribution

KJ formulated the central hypothesis of the study and drafted the manuscript; KJ, HP and LC conceived and designed the experiments; KJ, NI, JK, JRi, RR and LC performed the experiments; KJ, HP, HWH, RJC, PD, JK, JRo and LC analysed the data; HWH and RJC developed the mathematical models; KJ, HWH and RR contributed to the MS writing,

None of the authors have any competing interests

## Supporting Information

**Table S1. Allelic profiles of parents and their progeny within crossing experiments on the basis of seven microsatellite markers.**

**Table S2. Oocyte analysis.** For each female we indicate her biotype, origin, allozyme profile and the number of analyzed eggs.

**Table S3. Microsatellite markers and cytochrome** *b* **gene haplotypes.** This Table lists the alleles found in each sampled individual (specified by a respective ID number) on the basis of eight microsatellite markers. The cytochrome b gene haplotype is also specified.

**Appendix S4. SNP dataset.** Every contig is represented by ten lines, starting with its name (ref_acc) on the first line and followed by nine lines corresponding to nine individuals. Individuals’ genotypes are indicated by letters as follows: s, *C. strumicae;* p, *C. pontica;* e, *C. elongatoides;* t, *C. taenia;* n, *C. tanaitica*. Variant positions are expressed using IUPAC notation and invariant positions are excluded.

**Appendix S5. Summary of genetic divergences of fish species producing asexual hybrids.**

